# TMC1 and TMC2 are cholesterol-dependent scramblases that regulate membrane homeostasis in auditory hair cells

**DOI:** 10.1101/2025.07.03.663083

**Authors:** Hubert Lee, Yein Christina Park, Haosheng Wen, Harper E. Smith, Jayashree Balaraman, Runjia Cui, Marcos Sotomayor, Angela Ballesteros

**Affiliations:** Section on Sensory Physiology and Biophysics, National Institute on Deafness and other Communication Disorders, Bethesda, MD, USA; Graduate Program in Cell, Molecular, Developmental Biology & Biophysics, Johns Hopkins University, Baltimore, MD, USA; Biochemistry and Molecular Biophysics Graduate Program, University of Chicago, Chicago, IL, USA; Center for Mechanical Excitability, University of Chicago, Chicago, IL, USA; Department of Biochemistry and Molecular Biology, University of Chicago, Chicago, IL, USA; Biophysics Graduate Program, The Ohio State University, Columbus, OH, USA; Department of Biological Sciences, Rensselaer Polytechnic Institute, Troy, NY, USA

**Keywords:** Transmembrane channel-like proteins, TMC1 and TMC2, mechanosensitive ion channels, lipid scramblases, cholesterol, mechanoelectrical transduction, sensory hair cells, hearing and balance

## Abstract

TMC1 and TMC2, the pore-forming subunits of the mechanoelectrical transduction (MET) complex in inner ear sensory hair cells, are essential for auditory and vestibular function. Pathogenic mutations in TMC1 are a leading cause of genetic hearing loss, but their underlying cellular mechanisms remain poorly understood. Here, we reveal that TMC1 and TMC2 are cholesterol-regulated lipid scramblases whose activity modulates plasma membrane asymmetry. Using reconstituted proteoliposomes and molecular dynamics simulations, we demonstrate that both proteins facilitate phospholipid translocation across membrane bilayers, a process tuned by cholesterol and enhanced by deafness-causing TMC1 mutations. We show that this scramblase activity correlates with TMC1-dependent externalization of phosphatidylserine and membrane blebbing in murine auditory hair cells, linking TMC1-dependent membrane homeostasis dysregulation to auditory sensory cell pathology. These findings identify TMCs as a novel family of lipid scramblases, advancing our understanding of MET complex biology and offering mechanistic insight into membrane-driven forms of hereditary deafness.

## Introduction

Mechanosensitive (MS) channels are membrane proteins that respond to mechanical deformations of the lipid bilayer caused by a wide dynamic range of mechanical stimuli^1^. These channels are crucial for sensory processes such as hearing and balance, which rely on the fastest and most sensitive MS channels identified to date^2^. The MS channels in the inner ear, found in the auditory and vestibular sensory hair cells, are known as the mechanoelectrical transduction (MET) channels. These channels are located at the tips of the shorter rows of stereocilia, which are actin-rich protrusions of increasing heights arranged in precisely organized bundles^3^. Mechanical stimuli, such as sound waves (for auditory hair cells) or head movements (for vestibular hair cells), cause deflection of the stereocilia bundles towards the tallest stereocilium. This deflection causes tension in the extracellular protein filaments, known as tip links^4,5^, that connect adjacent stereocilia, ultimately leading to the opening of the MET channels and initiating the MET process that converts mechanical stimuli into electrical signals conveyed to the brain^6^.

After decades of investigation, the transmembrane channel-like (TMC) proteins 1 and 2 have been identified as the pore-forming subunits of the MET channel complex in auditory and vestibular inner ear hair cells and, accordingly, they have been reported to exhibit MS channel activity^7–9^. Recently, we showed that manipulations hindering MET function or decreasing intracellular Ca^2+^ led to phosphatidylserine (PS) externalization at the stereocilia bundle of murine auditory hair cells^10,11^. Importantly, this dysregulation of the hair cell membrane homeostasis does not occur in hair cells from *Tmc1^-/-^*;*Tmc2^-/-^* double knock out mice. Furthermore, mice carrying TMC1 deafness-causing missense mutations (*Mus musculus* MmTMC1 M412K, D569N, or D528N), known to diminish TMC1 MET channel function^12–15^, exhibit constitutively externalized PS^10,11,16^. Notably, auditory hair cells from all these TMC1 mutant mice degenerate and die, leading to hearing loss since mammalian hair cells do not regenerate^12,13,17^. These previous empirical observations and the evolutionary and structural relation of TMC proteins with TMEM16 and TMEM63/OSCA ion channels and lipid scramblases^7,18–21^ suggest that TMCs may function as lipid scramblases.

Like in most eukaryotic cells, membrane phospholipids are expected to be asymmetrically distributed in the phospholipid bilayer of the hair cell plasma membrane, with phosphatidylcholine (PC) and sphingolipids predominantly in the extracellular leaflet, and PS, phosphatidylethanolamine (PE), and phosphoinositide (PI) preferentially confined to the cytosolic leaflet. This lipid asymmetry is maintained by flippases and floppases, membrane proteins that translocate specific lipids against their concentration gradients in an ATP-dependent manner^22^. In contrast, lipid scramblases disrupt membrane asymmetry by facilitating energy-independent bidirectional lipid movement in accordance with lipid concentration gradients^22^. The activation of lipid scramblases leads to the externalization of anionic lipids, such as PS, which in turn alters the membrane properties and regulates essential physiological processes, including apoptotic cell clearance, blood coagulation, bone mineralization, and synaptic pruning^23^. In inner ear hair cells, the maintenance of membrane lipid asymmetry is critical for cell survival, as evidenced by the loss of hair cells and hearing in mice lacking or harboring mutations in various lipid flippases^24–27^ or, as mentioned above, mice carrying MET-disruptive mutations^11,16^. However, the mechanisms by which disruption of the auditory MET channels leads to constitutive PS externalization, resulting in hair cell degeneration and hearing loss, remain unknown.

Here, we show that the pore-forming subunits of the MET channel complex, TMC1 and TMC2, also function as lipid scramblases. Using well-established *in vitro* scramblase assays with purified proteins reconstituted into liposomes^28^, we assessed the potential TMC scramblase activity in a controlled membrane environment. Complementary molecular dynamics (MD) simulations^29^ we performed to gain insights into the lipid scrambling mechanisms. These *in vitro* an *in silico* findings were validated by functional assays in cochlear explants from wild type and three TMC1 mutant models of hearing loss. We demonstrate that TMC scramblase is finely tuned by cholesterol, and that three TMC1 deafness-causing mutations with enhanced scramblase activity led to disrupted plasma membrane asymmetry and PS externalization in auditory hair cells. Together, our findings identify TMCs a novel family of lipid scramblases and provide insights into the molecular mechanisms underlying hearing loss caused by disruption of cellular lipid homeostasis.

## Results

### *Chelonia mydas* TMC1 and *Melopsittacus undulatus* TMC2 function as lipid scramblases

Endogenous TMC1 and TMC2 localize to the plasma membrane of inner ear hair cell stereocilia^30^ but fail to traffic to the cellular membrane when expressed in heterologous systems^9,18,31^, hindering their functional characterization. Consequently, cell-based scramblase assays using the PS-binding protein annexin V, are not a viable approach to study the scramblase activity of wild-type TMC1 and TMC2. As shown in Supplemental Fig. S1, HEK293T cells transfected with the scramblase MmTMEM16F displayed obvious PS externalization on their plasma membrane while we did not detect annexin V staining in cells expressing the related chloride channel MmTMEM16A or in MmTMC1-transfected cells. To overcome this limitation, we decided to investigate the potential phospholipid scramblase (PLS) activity of TMC proteins using well-established *in vitro* liposome-based fluorescence assays^32,33^.

In these experiments, liposomes containing a small percentage of fluorescent 7-nitro-2,1,3-benzoxadiazol (NBD) acyl-labeled phospholipids are treated with sodium dithionite (SDT). SDT is a membrane-impermeable reducing agent that irreversibly quenches the NBD lipids accessible on the outer liposome leaflet, leading to a ∼50% reduction of the initial fluorescence. If a protein with PLS activity is incorporated into the liposomes, this activity will transport NBD lipids across leaflets making them accessible to SDT and resulting in >50% reduction of the initial fluorescence (Figure 1A). To control for the possibility that TMC proteins may form a pore through which SDT might gain access to NBD-labeled lipids in the inner leaflet of the membrane without being competent to scramble lipids, we also measured the reduction in NDB fluorescence produced by bovine serum albumin (BSA). BSA is a large protein that should not permeate the TMC pore but it can extract lipids from the outer leaflet and reduce their fluorescence^33^. Unlike SDT, BSA does not quench the fluorescence of NBD lipids but instead reduces it by half. Therefore, BSA reduces the total fluorescence in the liposome samples by a maximum of 25% or 50% in the absence or the presence of a scramblase, respectively (Figure 1B). In addition, proper protein reconstitution into liposomes was confirmed by SDS-PAGE (Figure 1C).

**Figure 1:**
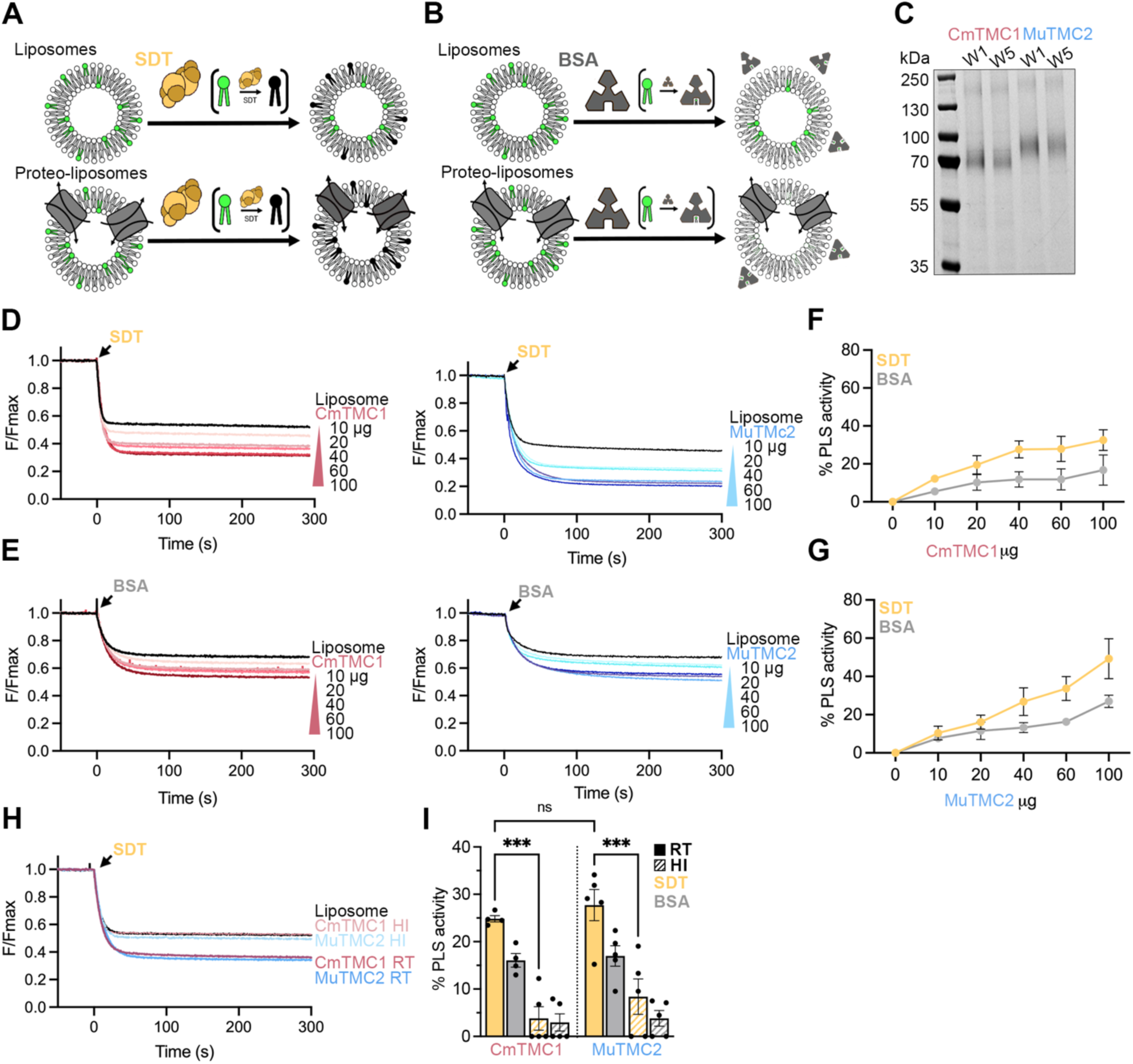
CmTMC1 and MuTMC2 are lipid scramblases. A. Cartoon representation of an SDT-based scramblase assay with liposome (top) or proteo-liposome (bottom) samples. B. Cartoon representation of an BSA-based scramblase assay with liposome or proteo-liposome samples. C. SDS-PAGE gel of CmTMC1 or MuTMC2 proteo-liposomes after the first (W1) or last (W5) BioBead washes showing the presence of protein in our samples. D. Decrease of NBD-fluorescence after SDT addition to liposomes (black) or proteo-liposomes containing increasing amounts of CmTMC1 (red) or MuTMC2 (blue). E. Decrease of NBD-fluorescence after BSA addition to liposomes (black) or proteo-liposomes containing increasing amounts of CmTMC1 (red) or MuTMC2 (blue). F. Quantification of CmTMC1 PLS activity in SDT-(yellow) and BSA-(grey) based assays. G. Quantification of MuTMC2 PLS activity in SDT-(yellow) and BSA-(grey) based assays. Mean ± SEM is shown for two to three independent experiments in F and G. H. Representative traces of NBD-fluorescence decrease after SDT addition in samples containing 60 μg of CmTMC1 (red) or MuTMC2 (blue) at room temperature (RT) or after heat inactivation (HI). I. SDT (yellow) and BSA (grey) PLS activity of CmTMC1 or MuTMC2 before (RT, solid) or after heat inactivation (HI, hollow). Mean ± SEM is shown. Each dot represents one independent experiment. Two-way ANOVA was performed to evaluate statistical significance. P values are indicated as: ns ≥ 0.05 and **** < 0.0001. See also Supplemental Fig. 3.

To test whether TMC proteins act as scramblases, we first focused on the truncated *Chelonia mydas* TMC1 (CmTMC1) and *Melopsittacus undulatus* TMC2 (MuTMC2) lacking the N- and C-terminal domains, which were identified among 21 TMC orthologs as amenable for protein expression and purification, and previously used to characterize their MS channel activity in liposome-based assays^8^. TMC proteins have 10 transmembrane segments (TM) and in some cases long intracellular N- and C-terminal domains^34,35^. A sequence alignment of these truncated CmTMC1 and MuTMC2 (MuTMC2Δ), and the database CmTMC1 (XP_043402179.1) and MuTMC2 (XP_033916589.1) (Supplemental Fig. 2), revealed that truncated MuTMC2Δ was also lacking the first extracellular loop, the transmembrane segment (TM) 2, and part of its TM10. Therefore, we expressed CmTMC1 containing residues 243-842, MuTMC2 containing residues 211-807, and the previously published MuTMC2Δ containing residues 210-775 (Supplemental Fig. 3A). Briefly, these proteins were expressed using the baculovirus expression system in Sf9 insect cells, solubilized in detergents, and purified by nickel affinity and size exclusion chromatography. Analysis by fluorescence detection size-exclusion chromatography (FSEC) showed that CmTMC1, MuTMC2, and MuTMC2Δ displayed minimal aggregation. We also confirmed that our proteins were of appropriate purity and had the expected size by SDS-PAGE (Supplemental Fig. 3B-E).

We next performed SDT- and BSA-based *in vitro* PLS assays with purified CmTMC1 and MuTMC2 reconstituted in liposomes containing palmitoyl-phosphocholine, -phosphoethanolamine, -phospho-L-serine, and cholesterol (POPC/PE/PS/CHO), which mimic the stereocilia lipid content^36^. To validate our experimental conditions, we first confirmed that purified *Homo sapiens* TMEM16K (HsTMEM16K), a Ca^2+^-dependent scramblase^37^, was able to scramble lipids in these liposomes in the presence of Ca^2+^ but not EGTA (Supplemental Fig. 3F-J).

As expected, SDT addition to POPC/PE/PS/CHO liposomes led to an approximate 50% reduction of the NBD fluorescence. However, SDT quenched >50% fluorescence in liposomes containing CmTMC1 or MuTMC2 in a protein dose-dependent manner (Figure 1D and F). Importantly, BSA control assays showed similar results (Figure 1E and G), supporting that the heightened NBD fluorescence reduction observed in the proteo-liposomes was due to scrambling of the fluorescent NBD-PE lipids by the TMC proteins and not caused by SDT leakage or permeation through the TMC pore. Furthermore, these samples lost their scramblase activity after heat-inactivation, underscoring the protein dependence of this effect (Figure 1H-I). Finally, in addition to NBD-PE, we also detected PLS activity in CmTMC1 and MuTMC2 proteo-liposomes containing a small percentage of NBD-labeled PG, PS, or PC, indicating that, like many other scramblases^32,37^, TMC proteins can translocate phospholipids with different headgroups (Supplemental Fig. 4). Therefore, we conclude that CmTMC1 and MuTMC2 are lipid scramblases with no specificity for the phospholipid headgroup.

### TMC lipid scramblase activity resides at the transmembrane core

Our previous data obtained using truncated CmTMC1 or MuTMC2 suggest that the TM core of the protein is sufficient to facilitate PLS activity. However, the presence of the intracellular N- and C-terminal domains might alter TMC lipid scrambling. Therefore, we next explored whether the physiological full-length (FL) proteins preserve scramblase function. To do this, we expressed and purified FL CmTMC1 and MuTMC2 using a similar expression and purification protocol. Compared to the truncated forms, these purified FL proteins displayed lower protein yields, some aggregates, and a wider FSEC peak, particularly for FL CmTMC1 (Supplemental Fig. 5A-D). However, SDT and BSA PLS assays showed no differences in PLS activity between truncated and FL CmTMC1 and MuTMC2, indicating that FL proteins function as lipid scramblases and confirming that the N- and C-terminal domains are not required for lipid scrambling (Supplemental Fig. 5E-H). Additionally, we measured the PLS activity of truncated MuTMC2Δ, which, in addition to the N- and C-terminal domains missing in MuTMC2, also lacks the first extracellular loop 1, the TM2, and part of the TM10 (Supplemental Fig. 2 and 3A). As shown in Supplemental Figure 5G-H, MuTMC2Δ, MuTMC2 and FL MuTMC2 exhibited similar PLS activity. Lastly, PLS assays with tagged and untagged CmTMC1 or MuTMC2 confirmed that the histidine and GFP tag did not affect lipid scrambling (Supplemental Fig. 5I-P). Altogether, these data indicate that the FL CmTMC1 and MuTMC2 proteins function as scramblases, that this activity resides at the TM core, and that it is not mainly mediated by the ECL1 or TM10.

### MET channel blockers and calcium have no direct effect on TMC scramblase activity

Well-established MET channel blockers such as the ototoxic aminoglycoside antibiotic neomycin, the diuretic benzamil, and the alkaloid curare have been shown to trigger PS externalization in mouse auditory hair cells in a TMC-dependent manner^11^. These data suggest that the channel and scramblase activities of TMC proteins may be inversely regulated. To test whether MET channel blockers could activate TMC scramblase activity, we first performed PLS assays with TMC proteo-liposomes generated in the absence or the presence of 0.1 mM of four chemically distinct MET blockers, tested independently. None of these treatments affected lipid scrambling by CmTMC1 or MuTMC2 (Figure 2A-D), indicating that the presence of MET blockers does not alter CmTMC1 or MuTMC2 scramblase activity *in vitro*.

**Figure 2:**
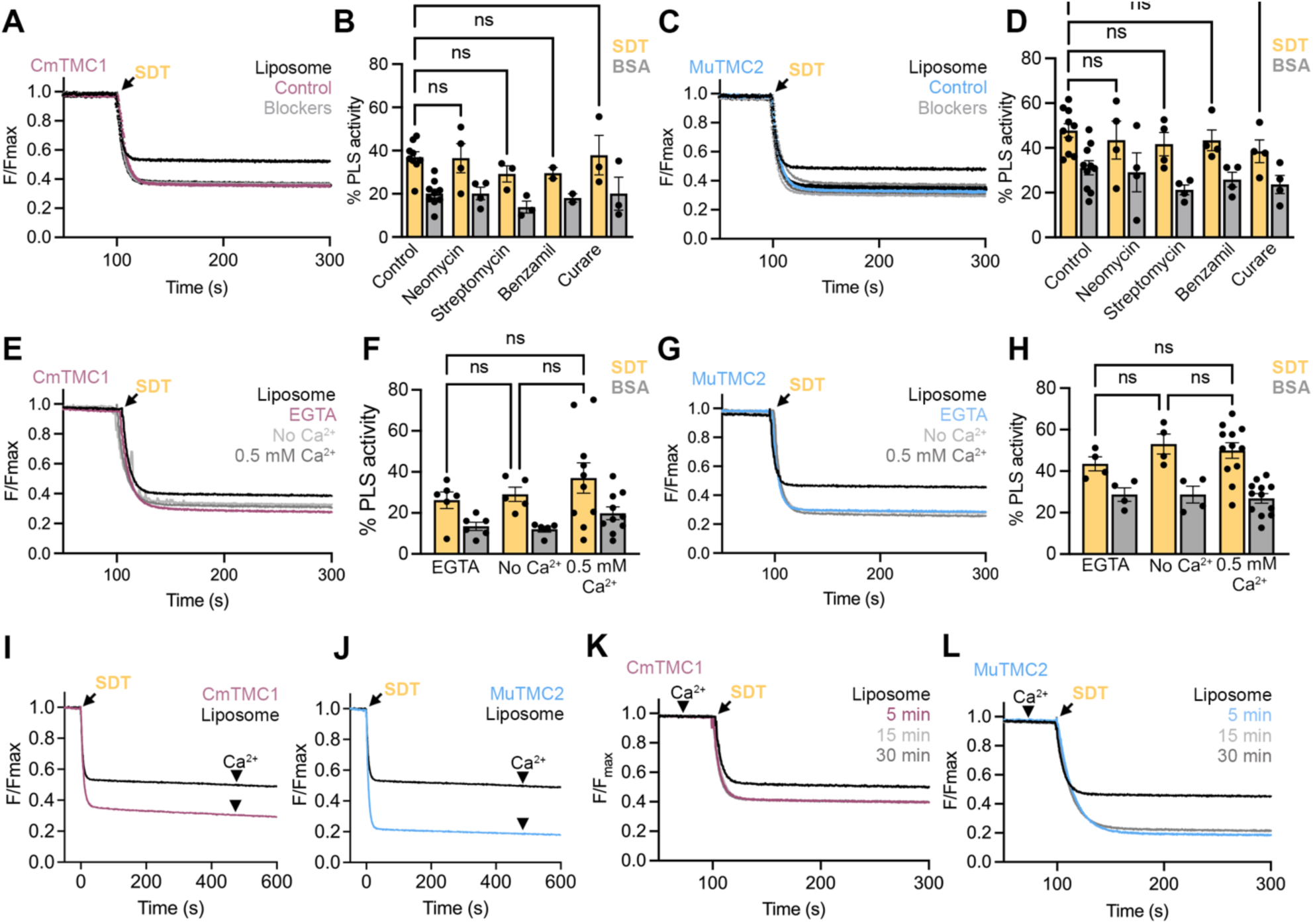
MET channel blockers and Ca^2+^ have no direct effect on TMC scramblase activity *in vitro*. A. Representative traces for NBD fluorescence decrease due to SDT addition (arrow) to liposomes (black) or proteo-liposomes containing CmTMC1 in buffer (control) or in the presence of 0.1 mM neomycin, streptomycin, benzamil, or curare (grey). B. Quantification of CmTMC1 SDT-(yellow) and BSA-(grey) based PLS activity in the conditions shown in A. C. Similar data to that shown in A for proteo-liposomes containing MuTMC2. D. Quantification of MuTMC2 PLS activity in the conditions shown in C. E. Representative traces of NBD fluorescence decrease due to SDT addition in liposomes (black) or CmTMC1 proteo-liposomes in the absence (light grey), or presence of EGTA (red) or Ca^2+^ (dark grey). F. Quantification of SDT-(yellow) and BSA-(grey) based CmTMC1 PLS activity in the conditions shown in E. G. Representative traces of NBD fluorescence decrease due to SDT addition (arrow) in liposomes (black) or MuTMC2 proteo-liposomes in the absence (light grey) or presence of EGTA (red) or Ca^2+^ (dark grey). H. Quantification of SDT-(yellow) and BSA-(grey) based MuTMC2 PLS activity in the conditions shown in G.I-J. Representative traces of NBD fluorescence decrease due to SDT (arrow) and Ca^2+^ (arrowhead) addition to liposomes (black) or CmTMC1(I) or MuTMC2 (J) proteo-liposomes. K-L. Representative traces of NBD fluorescence decrease due to SDT (arrow) addition to liposomes (black) or CmTMC1 (K) or MuTMC2 (L) proteo-liposomes pre-incubated with Ca^2+^ for 5, 10, or 30 min. Mean ± SEM is shown for B, D, F and H. Each dot represents one independent experiment. One-way ANOVA was performed to evaluate statistical significance. P values are indicated as: ns≥0.05.

Blockade of the MET channel by open-channel pore blockers may lead to a reduction of intracellular Ca^2+^ in hair cells since cations are not able to permeate through an obstructed MET channel. Intracellular Ca^2+^ is known to regulate the PLS activity of TMEM16 lipid scramblases^38,39^. In auditory hair cells, intracellular Ca^2+^ levels have been inversely correlated with PS externalization^40^. Consistent with this, treatment of auditory hair cells from wild-type mice with the intracellular Ca^2+^ chelator BAPTA-AM triggered externalization of PS^11^. Therefore, MET blockers may play an indirect role in PS externalization by reducing the intracellular Ca^2+^ concentration. Therefore, to understand the effect of Ca^2+^ in TMC-mediated lipid scrambling, we analyzed the PLS activity in CmTMC1 and MuTMC2 proteo-liposomes prepared in the presence of Ca^2+^ or EGTA or in the absence of both, where the extra- and intraliposomal solutions present similar composition. We observed comparable levels of lipid scrambling by CmTMC1 and MuTMC2 in these three conditions (Figure 2E-H), indicating that, unlike TMEM16s^32,37^ (Supplemental Fig. 3H), TMC-mediated lipid scrambling is not directly regulated by Ca^2+^. Similarly, we did not detect changes in the CmTMC1 and MuTMC2 PLS activity when Ca^2+^ was added after SDT addition to proteo-liposomes generated in the absence of Ca^2+^, when the extraliposomal Ca^2+^ concentration is set to be temporarily higher than the intraliposomal (Figure 2I-J), or when we pre-equilibrated the liposomes with Ca^2+^ for different times (5, 15, and 30 min) before measuring the NBD fluorescence (Figure 2K-L). Together, these results indicate that MET blockers and Ca^2+^ do not directly regulate lipid scrambling by CmTMC1 or MuTMC2 *in vitro*.

### Cholesterol regulates TMC1 and TMC2 lipid scramblase activity

MS channels are modulated by lipids through direct interactions or, indirectly, by altering the physical properties of the membrane^41^. Consistently, the properties of the inner ear MET channel are modulated by the lipid enviroment^42,43^, suggesting that lipids regulate TMC1 and TMC2 MS channel function. Additionally, recent studies have shown that cholesterol regulates the PLS activity of opsin and TMEM63B^20,44^, implying a role for the membrane in regulating lipid scramblase activity. Considering these observations, we next investigated the effect of cholesterol and the lipid environment on CmTMC1 and MuTMC2 scrambling.

We first analyzed whether cholesterol is required for TMC lipid scrambling *in vitro* by reconstituting CmTMC1 or MuTMC2 in POPC/PE/PS liposomes without (0%) or with 10% cholesterol (Figure 3). While performing these assays, we realized that proteins purified in DDM detergent containing cholesterol (DDM/CHS) may contain some residual cholesterol from the solubilization and purification steps. Therefore, we also performed PLS assays with proteins purified in Fos-Choline 12 (FC12) detergent (Supplemental Fig. 6A-D). Proteins purified in FC12 presented slightly lower activity than those purified in DDM/CHS, regardless of the liposomal cholesterol content. However, under both purification conditions, we found a reduction in the PLS activity of CmTMC1 and MuTMC2 in proteo-liposomes lacking cholesterol compared to those containing 10% cholesterol, with MuTMC2 exhibiting higher basal PLS activity than CmTMC1 in the absence of cholesterol (Figure 3A-C). We next quantified the cholesterol content in our samples to estimate the amount of cholesterol required to trigger TMC PLS activity. This analysis showed that CmTMC1 and MuTMC2 require 8 1 μg/mL and 136 μg/mL of cholesterol, respectively, to reach half of their maximum scramblase activity. These results suggest that cholesterol can enhance TMC scrambling and that CmTMC1 and MuTMC2 PLS activities have different cholesterol requirements.

**Figure 3:**
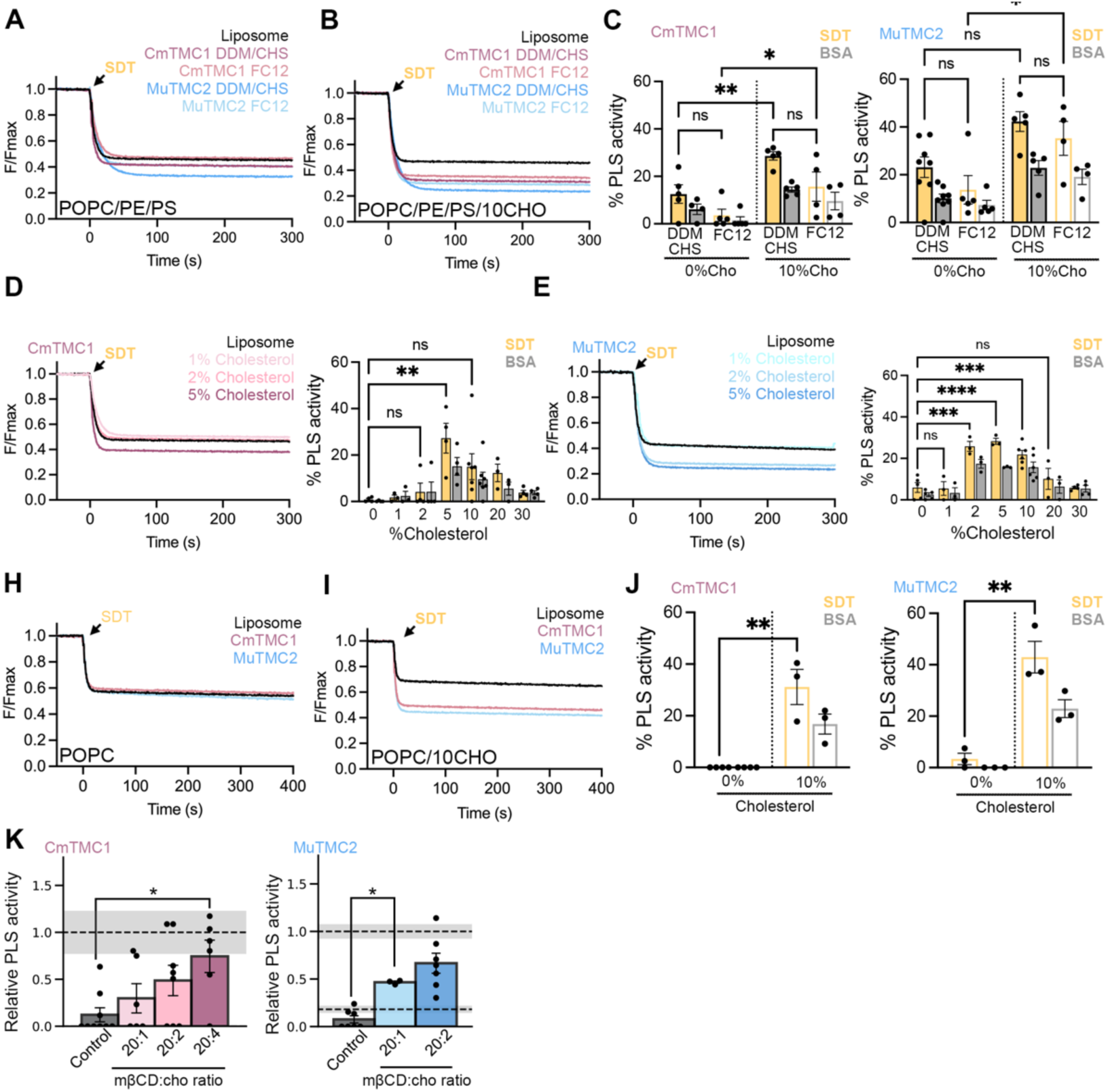
Cholesterol promotes TMC1 and TMC2 scramblase activity. A. NBD-fluorescence decrease after SDT addition to POPC/PE/PS liposomes (black) or proteo-liposomes containing CmTMC1 purified in DDM/CHS (red) or FC12 (pink) or MuTMC2 purified in DDM/CHS (blue) or FC12 (light blue). B. NBD-fluorescence decrease after SDT addition to POPC/PE/PS liposomes containing 10% cholesterol (black) or proteo-liposomes containing CmTMC1 purified in DDM/CHS (red) or FC12 (pink) or MuTMC2 purified in DDM/CHS (blue) or FC12 (light blue). C. Quantification of the SDT-(yellow) and BSA-(grey) PLS activity of CmTMC1 (left) or MuTMC2 (right) purified in the conditions represented in A and B. D. NBD-fluorescence decrease after SDT addition to POPC/PE/PS liposomes (black) or CmTMC1 proteo-liposomes (red) containing increasing amounts of cholesterol (left) and quantification of CmTMC1 PLS activity in these assays (right). E. Decrease of NBD-fluorescence after SDT addition to POPC/PE/PS liposomes (black) or MuTMC2 proteo-liposomes (blue) containing increasing amounts of cholesterol (left) and quantification of MuTMC2 PLS activity in these conditions (right). H. NBD-fluorescence decrease after SDT addition to POPC liposomes (black) or proteo-liposomes containing CmTMC1 (red) or MuTMC2 (blue). I. Decrease of NBD-fluorescence after SDT addition to POPC liposomes containing 10% cholesterol (black) or proteo-liposomes containing CmTMC1 (red) or TMC2 (blue). J. Quantification of the SDT-(yellow) and BSA-based (grey) PLS activity of CmTMC1 (left) or MuTMC2 (right) of the representative data shown in H and I. K. CmTMC1 (left) or MuTMC2 (right) PLS activity, normalized by average PLS activity at 10% cholesterol, after cholesterol addition via treatment with empty mβCD (control) or mβCD/CHO complexes with increasing cholesterol amounts. Grey bars indicate PLS activity at 0% (bottom) or 10% (top) cholesterol. Mean ± SEM is shown for C, D, E, J and K. Each dot represents one independent experiment. Two (C) or one-way ANOVA (D-E, J-K) was performed to evaluate statistical significance. P values are indicated as: ns≥0.05, *<0.05, **<0.01, ***<0.001 and ****<0.0001. See also Supplemental Fig. 6A-D.

To further explore the role of cholesterol in TMC scramblase activity, we performed additional assays with POPC/PE/PS liposomes containing increasing amounts of cholesterol: 0%, 1%, 2%, 5%, 10%, 20%, and 30%. These dose-response assays revealed a biphasic response where both low (<2-5%) and high (>20%) cholesterol levels hinder TMC PLS activity (Figure 3D-E). Additionally, the PLS activity dependence on cholesterol differed for CmTMC1 and MuTMC2, with CmTMC1 reaching maximal activity in liposomes prepared with 5% cholesterol, whereas MuTMC2 displayed maximal activity in liposomes prepared with 2-10% cholesterol. Interestingly, like opsin and TMEM63B^20,44^, CmTMC1 and MuTMC2 showed little to no activity in proteo-liposomes containing ≥30% cholesterol, suggesting that inhibition of PLS activity by high levels of cholesterol may be a conserved feature of lipid scramblases to limit unchecked lipid scrambling at the mammalian plasma membrane.

To rule out the role of POPE and POPS in TMC scramblase activity, we next performed PLS experiments with liposomes containing only POPC without or with 10% cholesterol. As shown in Figure 3H-J, like in POPC/PE/PS liposomes (Figure 3C), POPC proteo-liposomes containing CmTMC1 or MuTMC2 purified in FC12 required cholesterol for lipid scrambling. Furthermore, we were able to activate TMC-mediated scrambling in cholesterol-free POPC proteo-liposomes by supplementing cholesterol using cyclodextrin and cholesterol (mβCD/CHO) complexes of increasing cyclodextrin:cholesterol ratios, which are commonly used to donate cholesterol to liposomes and cells^44,45^. On the other hand, incubation of POPC proteo-liposomes with mβCD lacking cholesterol (control) did not increase the PLS activity of CmTMC1 and MuTMC2, further supporting the role of cholesterol in TMC lipid scrambling. Consistent with the specific cholesterol requirements of CmTMC1 and MuTMC2, we detected MuTMC2 scramblase activity when proteo-liposomes were incubated with 20:1 mβCD/CHO complexes, while CmTMC1 PLS activity required 20:2 mβCD/CHO complexes containing more cholesterol (Figure 3K). Importantly, since cholesterol was added after protein incorporation into the liposomes, these experiments ruled out potential artifacts arising from variations in protein reconstitution efficiency across liposomes with different cholesterol percentage (Figure 3D-E). These data highlight the role of cholesterol as the main lipid regulator of CmTMC1 or MuTMC2 PLS activity.

### Molecular dynamics simulations support cholesterol-dependent TMC lipid scrambling

To gain a molecular-level understanding of TMC-mediated lipid translocation across the bilayer leaflets, we prepared *in silico* TMC systems and performed molecular dynamics (MD) simulations^29^. We first predicted dimeric AlphaFold3^46^ structural models of truncated CmTMC1 and MuTMC2, which had a topology consistent with previous experimentally validated models of mouse and human TMC1 and TMC2 proteins^7,18,47,48^. These structural models were then embedded in POPC/PE/PS bilayers, either without or with 10% cholesterol (Supplemental Fig. 7A and Supplemental Table 1). Initial 100-ns long equilibrations were followed by targeted molecular dynamics (TMD) simulations opening one of the subunits of the dimers^49,50^ (Supplemental Fig. 7B-C). The TMD simulations were guided by a previously generated homology model of MmTMC1 based on open structures of TMEM16 proteins^18^. After the TMD simulations, each dimeric system was simulated for up to 1.7 μs with positional constraints to keep the open-like pore conformation. In each system, the other subunit of the dimer was kept in a closed conformation serving as a control for scramblase activity.

Analysis of the all-atom MD trajectories revealed different lipids approaching and interacting with the hydrated TMC open-pore groove. We observed three complete scrambling events through the open pores of the CmTMC1 and MuTMC2 systems with cholesterol. A PE lipid scrambled from the extracellular to the cytoplasmic side through CmTMC1, while one PC and one PE lipid scrambled together from the intracellular to the extracellular side through MuTMC2 (Figure 4A, Supplemental Figure 7D, and Movies 1 and 2). These results support that TMC proteins can scramble lipids.

**Figure 4:**
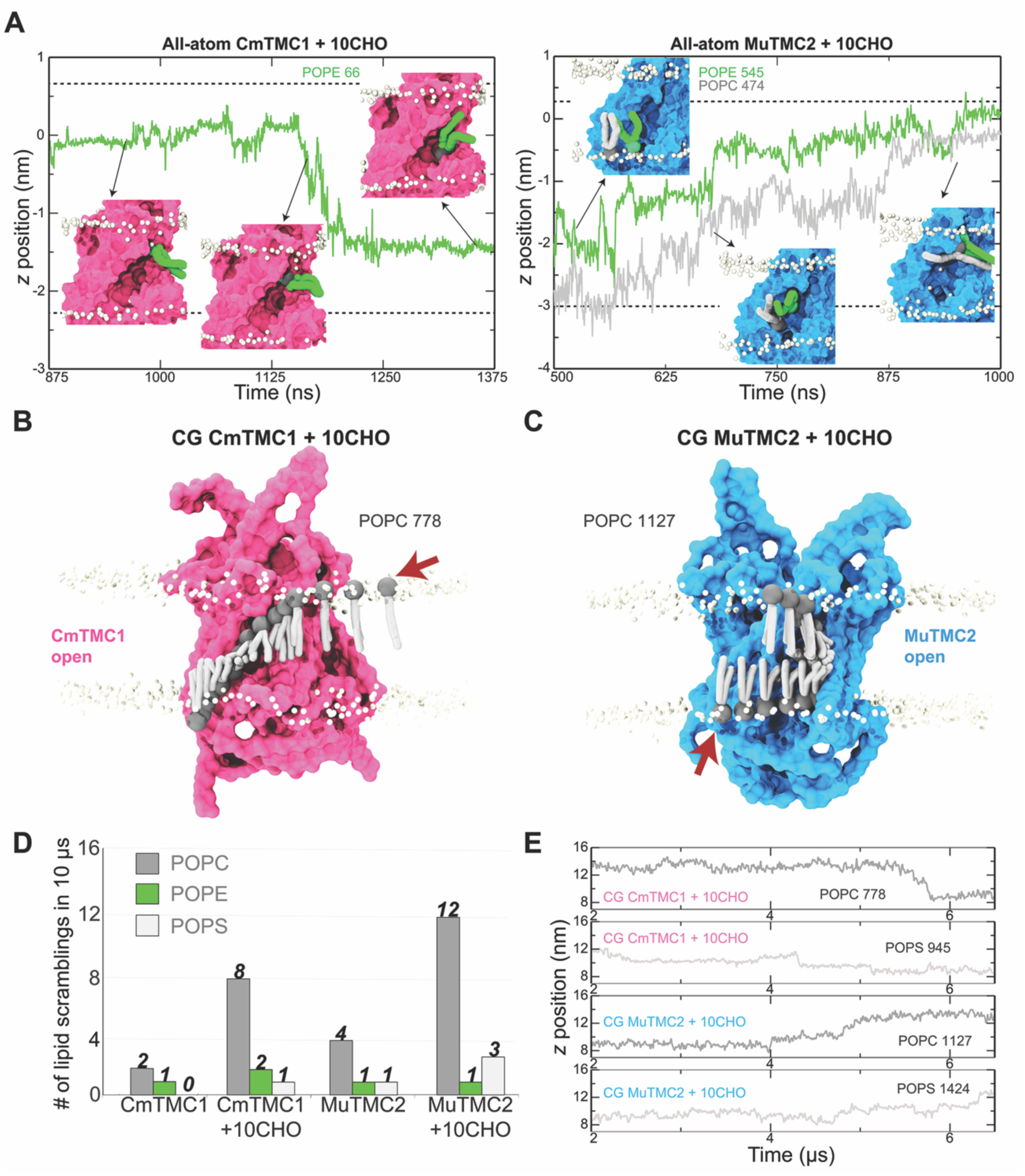
MD simulations support cholesterol-dependent TMC lipid scrambling at the canonical pathway. A. Plot of head-group position perpendicular to the membrane plane (*z*) as a function of time for lipids (POPC in gray and POPE in green) that translocated through the open grooves of CmTMC1 (left) and MuTMC2 (right) during equilibrium all-atom MD simulations with 10% cholesterol. Insets are snapshots at indicated time points. Protein is shown in molecular surface, non-translocating lipid phosphate atoms are white spheres. Water and ions omitted for visualization purposes. B-C. Canonical lipid scrambling pathway in the TMD-induced open pore of CmTMC1 (B) and MuTMC2 (C) during CG simulations with cholesterol. In each case a single lipid is shown at different time points overlaid on a representative structural model of the protein. Non-translocating lipid phosphate atoms are shown as white spheres. Red arrows indicate starting position in trajectory. D. Summary of scrambling events detected in 10-μs long CG simulations of CmTMC1 and MuTMC2 systems without and with 10% cholesterol. Each bar represents the total number of lipid translocation events by lipid type (POPC, POPE, and POPS). E. Lipid head-group *z*-position perpendicular to the membrane as a function of time for POPC (dark gray) and POPS (light gray). These lipids translocated through the open-like pores of CmTMC1 and MuTMC2 during CG MD simulation with cholesterol. *See also Supplemental* Fig. 7.

The limited number of scrambling events reflects the intrinsic timescale limitations of all-atom MD. To overcome this limitation, we conducted longer coarse-grained (CG) MD simulations. Analysis of 10 μs-long CG trajectories revealed non-selective CmTMC1 and MuTMC2 lipid scrambling with a strong cholesterol dependence (Figure 4B-E), reinforcing our *in vitro* experimental data. Simulations of the CmTMC1 system without cholesterol revealed 2 PC and 1 PE scrambling events, corresponding to a scrambling rate of approximately 3×10^5^ lipids·s^-1^, while 11 events (8 PC, 2 PE, and 1 PS) were observed in the presence of cholesterol, yielding a higher scrambling rate of 1.1×10^6^ lipids·s^-1^. Similarly, we observed 6 events (4 PC, 1 PE, and 1 PS) for the MuTMC2 system without cholesterol (6×10^5^ lipids·s^-1^) and 16 events (12 PC, 1 PE, and 3 PS) in the presence of cholesterol (1.6×10^6^ lipids·s^-1^) (Figure 4B-E and Supplemental Table 1). CG simulations are known to overestimate scrambling rates^51,52^, which can be up to 10^5^ lipids·s^-1^ or faster in experiments^53^. Nevertheless, our results suggest that TMC1 and TMC2 can function as highly efficient, cholesterol-regulated lipid scramblases, positioning TMC proteins as effective lipid translocators.

Last, we analyzed the predicted lipid translocation pathways to gain insight on whether TMC scrambling occurs at the canonical site, the TM4-TM6 pore^53^, or through the non-canonical site located at the TM10 dimerization region^51,54^. This analysis revealed that almost all events (33 out of 36) occurred through the canonical open TM4-TM6 groove across all four TMC systems (Movie 3). Unexpectedly, one PC lipid translocated through the closed CmTMC1 pore in the absence of cholesterol (Supplemental Figure 7E and Movie 4), and one PC and one PS scrambled through the noncanonical dimerization pathway in the MuTMC2 system containing cholesterol (Supplemental Figure 7E and Movies 4 and 5). These findings show that while closed pore and non-canonical translocation routes exist, the TM4-TM6 open pore serves as the primary pathway for TMC lipid scrambling.

### Cholesterol influences TMC lipid scrambling *in vitro* and in murine auditory hair cells

We next analyzed whether MmTMC1 and MmTMC2 intrinsically function as lipid scramblases. Using a similar protocol to the one implemented for CmTMC1 and MuTMC2, we purified MmTMC1 and MmTMC2 (Supplemental Fig. 6E-F) and reconstituted them in POPC/PE/PS liposomes without or with 10%, or 30% cholesterol, which resembles the physiological cholesterol levels^36^. Consistent with our previous results, we found that MmTMC1 and MmTMC2 scramble lipids in the presence of 10% cholesterol but not in liposomes lacking cholesterol or containing 30% cholesterol (Figure 5A-D). These results demonstrate that MmTMC1 and MmTMC2, like CmTMC1 and MuTMC2, function as cholesterol-dependent lipid scramblases, indicating that lipid scrambling is a conserved feature of TMC1 and TMC2 proteins.

**Figure 5:**
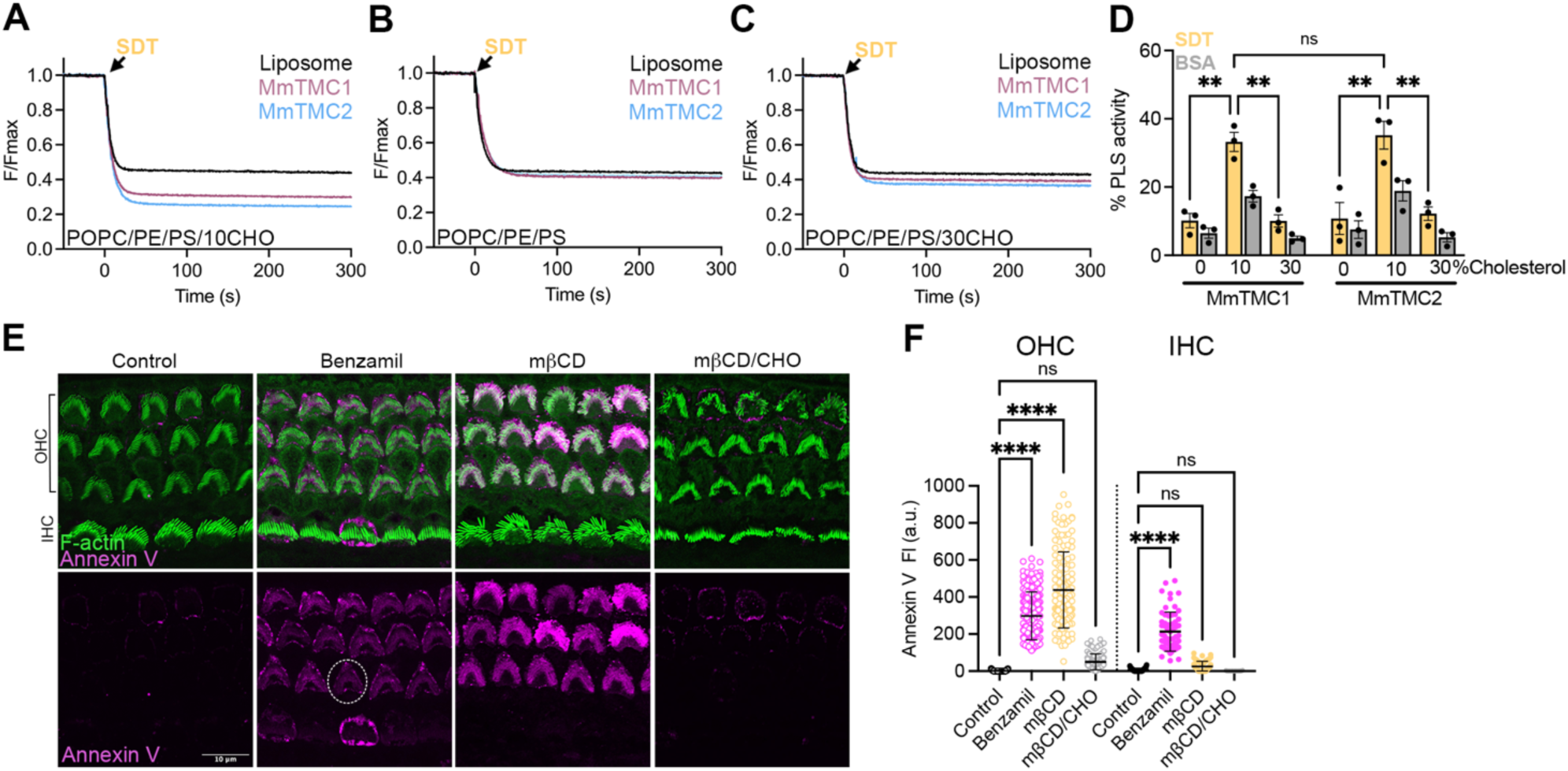
Cholesterol levels influence mouse TMC1 and TMC2 scramblase activity. NBD-fluorescence decrease after SDT addition to POPC/PE/PS liposomes (black) or MmTMC1 (red), or MmTMC2 (blue) proteo-liposomes containing 10% (A), 0% (B), or 30% cholesterol (C). D. Quantification of the MmTMC1 and MmTMC2 SDT-(yellow) and BSA-based (grey) PLS activity in the absence or presence of 10% and 30% cholesterol. Mean ± SEM is shown, and each dot represents one independent experiment. Two-way ANOVA was performed to evaluate statistical significance. E. Representative confocal images of auditory OHCs and IHCs at the middle cochlear region from P6 wild-type mice incubated in HBSS in the absence (control) or presence of benzamil, mβCD, or mβCD loaded with cholesterol (mβCD/CHO) and labeled with annexin V (magenta) and spy-555 (green) to visualize PS and F-actin, respectively. F. Quantification of background-subtracted annexin V-647 fluorescence intensity from the experimental conditions represented in E. Each dot represents the annexin V fluorescence intensity from one hair cell. Mean ± SEM is shown for n ∼ 100 OHCs and 30 IHCs from 3 tissues. One-way ANOVA was performed to evaluate statistical significance between control and treated samples. P values are indicated as: ns ≥ 0.05, ** < 0.01, and **** < 0.0001. See also Supplemental Fig. 6E-F.

Next, we explored the role of cholesterol in PS externalization in mouse auditory hair cells. As shown in Figure 5E-F, and consistent with previously published data^10,11^, the MET blocker benzamil triggered PS externalization in outer and inner auditory hair cells (OHC and IHC, respectively). Interestingly, treatment of cochleae tissue with mβCD/CHO complexes led to a minimal increase in annexin V signal while treatment with mβCD alone, which reduces membrane cholesterol levels^45^, triggered PS externalization in OHCs (Figure 5E-F). Cholesterol quantification in these conditions confirmed the efficacy of these treatments and revealed altered cholesterol levels from 4.70 ± 0.15 (n = 4) in untreated, to 2.81 ± 0.34 µg/mL (n = 4) in mβCD-treated cochleae, and 7.23 ± 0.21 µg/mL (n = 3) in mβCD/CHO-treated samples. These results indicate that MmTMC1 and MmTMC2 act as lipid scramblases regulated by cholesterol both *in vitro* and *in vivo*.

### Deafness-causing mutations in TMC1 enhance its scramblase activity

Hair cells from mice carrying the dominant M412K, D569N, or recessive D528N deafness-associated mutations (*Tmc1*^M412K/M412K^, *Tmc1*^D56N/D569N^, or *Tmc1*^D528N/D528N^), located at the putative TMC1 pore region (Figure 6A), exhibit constitutive externalization of PS at the apical hair cell region^10,11,16^ and profound progressive hearing loss^12,13,17^. We hypothesized that PS externalization in these mice may be caused by enhancement or dysregulation of the TMC1 scramblase function. To test this, we expressed and purified the equivalent CmTMC1 mutants (M507K, D658N, and D617N) (Supplemental Fig. 6G-H) and studied their scramblase activity in POPC/PE/PS liposomes without and with 10% or 30% cholesterol. In cholesterol-free proteo-liposomes with comparable amounts of protein, wild-type CmTMC1 and all three TMC1 mutants displayed similar levels of PLS activity (Figure 6B and E), and consistent with our previous results, the scramblase activity of all proteins was lower in the absence (0%) or presence of 30% liposomal cholesterol than in liposomes with 10% cholesterol (Figure 6B-E). Interestingly, the dominant CmTMC1 M507K and D658N deafness-causing mutants exhibited slightly higher PLS activity than the wild-type protein in liposomes containing 10% cholesterol (Figure 6C and G), and all three mutants tested exhibited enhanced PLS activity in the presence of 30% cholesterol (Figure 6D-E and H). These results indicate that the deafness-causing mutations enhance TMC1 PLS activity at physiological cholesterol levels. We also observed no differences in the kinetics of the NBD fluorescence decay rates after SDT addition between wild type and mutants CmTMC1 at any of the cholesterol percentages tested (Figure 6B-D), suggesting that the differences in PLS activity between these proteins is a consequence of the mutations endowing CmTMC1 scramblase activity at wider cholesterol ranges^44^. Importantly, depletion of cholesterol in auditory hair cells from *Tmc1*^M412K/M412K^ or *Tmc1*^D569N/D569N^ mice by mβCD treatment, which we confirmed decreased the cochleae cholesterol levels from 4.79 ± 0.17 to 2.85 ± 0.06 µg/mL (n = 4), significantly reduced the levels of constitutively externalized PS in OHCs (Figure 6I-L). These data support that constitutive externalization of PS observed in hair cells from mice carrying TMC1 deafness-causing mutations is caused by enhancement of the TMC1 lipid scramblase activity at physiological cholesterol levels. All together, we conclude that TMC1 and TMC2 are scramblases that regulate hair cell membrane homeostasis by their cholesterol-dependent lipid scramblase activity.

**Figure 6:**
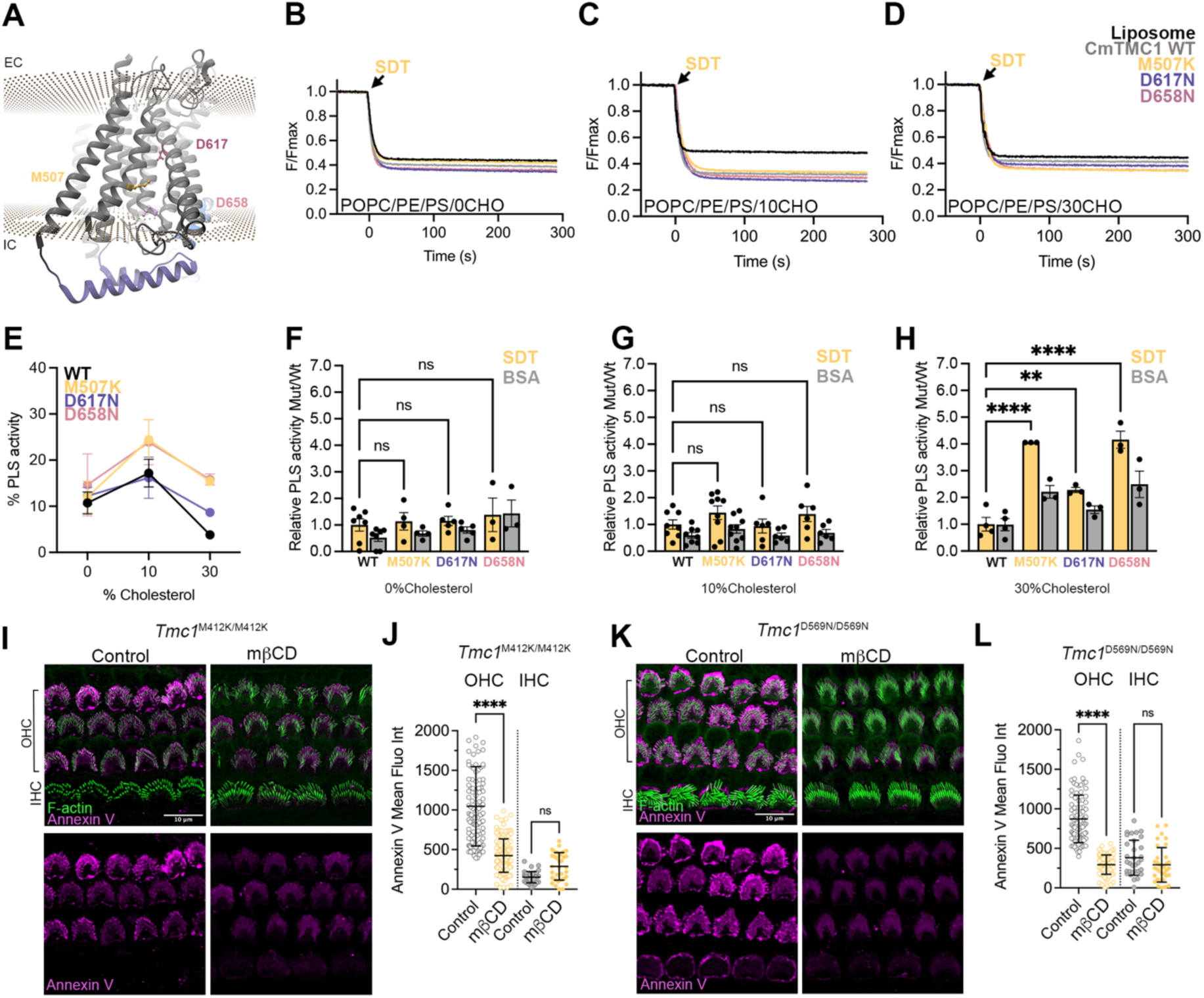
Deafness-causing mutations in TMC1 increase scramblase activity by altering its dependence on cholesterol. A. Localization of the equivalent residues to mouse TMC1 M412 (M507), D528 (D617) and D569 (D658) in the *Caenorhabditis elegans* TMC1 structure (PDB ID: 7USX). The approximate localization of the plasma membrane as calculated by the PPM web server is indicated by two parallel planes of spheres. NBD-fluorescence decrease after SDT addition to POPC/PE/PS liposomes (black) or proteo-liposomes containing wild type (gray) or mutants CmTMC1 M507K (yellow), D617N (purple) or D658N (pink) in liposomes containing 0% (B), 10% (C), or 30% (D) cholesterol. E. Quantification of the SDT-based PLS activity of wild-type CmTMC1, M507K, D617N or D658N mutants in POPC/PE/PS liposomes containing 0, 10 or 30 % cholesterol. F-H. SDT (yellow)- and BSA (grey)-based PLS activity of M507K, D617N and D658N mutants relative to wild-type CmTMC1 in proteo-liposomes containing 0% (F), 10% (G) or 30% (H) cholesterol. One-way ANOVA was performed to evaluate statistical significance between wild type and mutant TMC1 relative SDT-based activity. I-L. Representative confocal images of auditory OHCs and IHCs at the middle cochlear region from Tmc1^M412K/M412K^ (I) or Tmc1^D569N/D569N^ (K) mice incubated in HBSS without (control) or with mβCD to remove cholesterol and labeled with annexin V (magenta) or spy-555 (green) to visualize PS or F-actin. J and L. Quantification of background-subtracted annexin V-647 fluorescence intensity from the experiments showed in I or K. Mean ± SEM is shown for n∼ 100 OHCs and 30 IHCs from 3 tissues. Two-way ANOVA was performed to evaluate statistical significance between control and mβCD-treated samples in IHCs or OHCs. P values are indicated as: ns≥0.05, ** <0.01, *** <0.001, and ****<0.0001.See also Supplemental Fig. 3G-H.

## Discussion

In this study, we demonstrate that TMC1 and TMC2 are cholesterol-regulated lipid scramblases. Using *in vitro* liposome-based PLS assays, we show that CmTMC1, MuTMC2, MmTMC1 and MmTMC2 scramble a variety of phospholipids across lipid bilayers. This PLS activity is modulated by cholesterol, with high (> 10%) or low (< 2%) levels hampering the *in vitro* PLS activity of TMC1 and TMC2 (Figure 7A). MD simulations confirm the cholesterol dependance of the TMCs PLS activity and point to the TM4-TM6 groove as the main lipid translocation pathway. In addition, we show that the cholesterol range for optimal TMC1 PLS activity is widened in three different hearing loss-associated TMC1 mutations, enabling these mutant TMC1 proteins to scramble lipids at physiological levels of cholesterol. Consistent with this and confirming previous data from our lab, we demonstrate that untreated auditory hair cells from TMC1 mutant mice, but not wild-type mice, exhibit constitutive PS externalization. Furthermore, we show that reduction of the plasma membrane cholesterol levels by mβCD treatment of auditory hair cells activates PS externalization in wild-type hair cells, which supports a role for cholesterol in the regulation of the TMC1 PLS activity *in vivo*. Previous studies have supported the existence of a TMC-dependent hearing loss mechanism associated with the dysregulation of hair cell plasma membrane homeostasis^10,11,16^. Our data identify the mechanistic basis for this membrane-related hearing loss and indicate that TMC1 and TMC2 can function as both MS channel and lipid scramblases.

**Figure 7:**
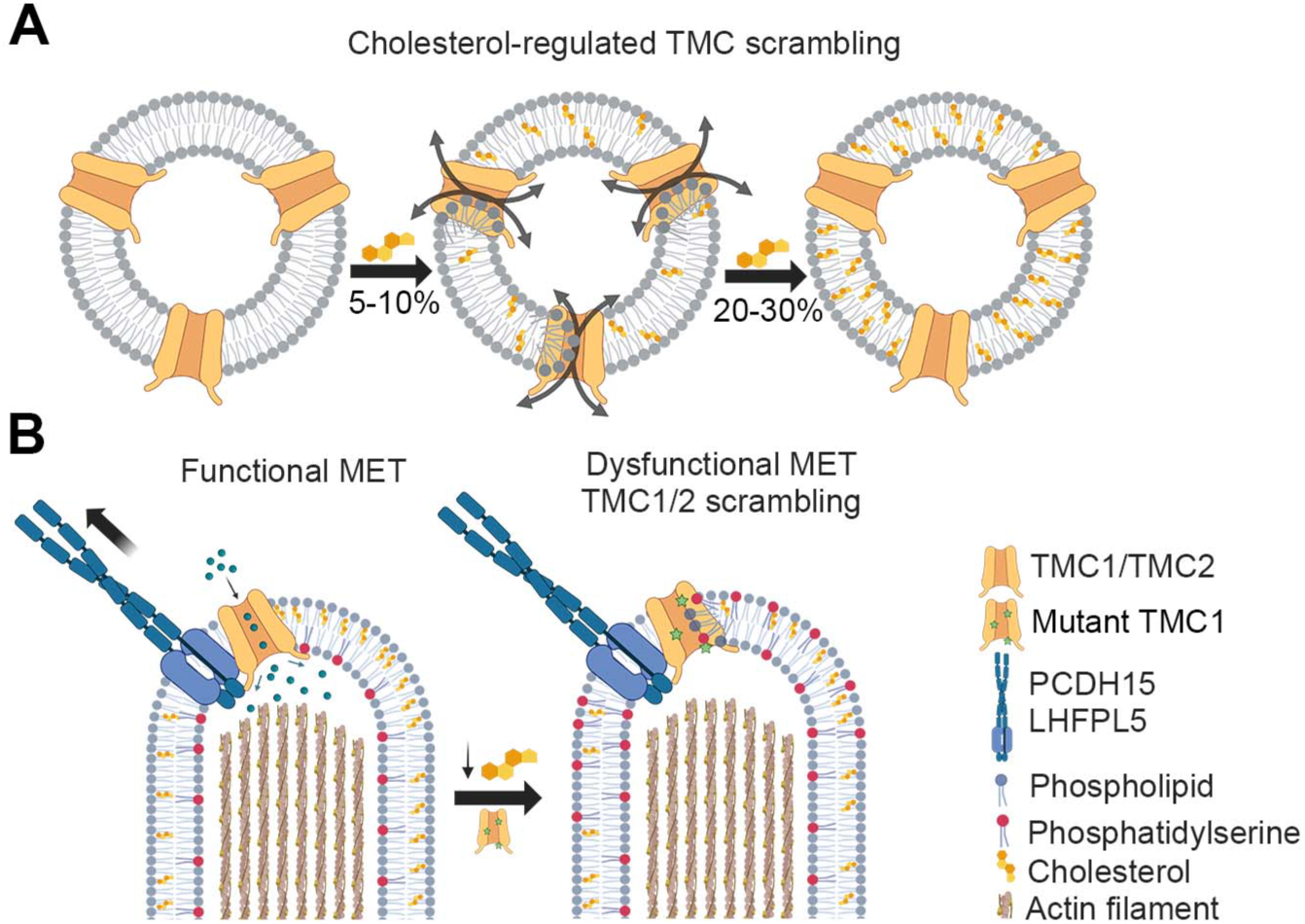
Cholesterol regulates TMC1 and TMC2 scramblase activity. A. *In vitro* scramblase assays with purified proteins reconstituted into liposomes containing varying cholesterol concentrations showed that TMC1 and TMC2 scramblase activity is maximal at 2-5% cholesterol but is inhibited at higher or lower cholesterol levels. B. TMC scramblase function is also regulated by cholesterol *in vivo*. Manipulations that alter cholesterol levels in auditory hair cell explants from wild type or two TMC1 deafness-causing mutant mice alter PS externalization.

Previous experiments in murine auditory hair cells showed that genetic, chemical, or mechanical disruption of the MET process leads to dysregulation of membrane homeostasis, manifested in loss of the plasma membrane asymmetry, PS externalization^10,16,55^, and membrane blebbing that results in the removal of TMC1 from the hair cell membrane^11^. Because these manipulations hindered ion permeation through the MET channels, and intracellular Ca^2+^ levels were found to inversely correlate with PS externalization^11,40^, previous studies pointed to Ca^2+^ as the main regulator for hair cell membrane homeostasis. However, our data shows that TMC scramblase activity *in vitro* is not affected by MET blockers or Ca^2+^. It is worth noting that in our experiments, TMCs may present a closed pore that impedes blocker binding due to the absence of the resting tension that the MET channel complex experiences in physiological conditions^4,56^. Furthermore, TMCs, like TMEM63/OSCA, lack the Ca^2+^-binding site conserved in the Ca^2+^-activated TMEM16 scramblases^57,58^, hinting at a distinct regulatory mechanism. Supporting this, we demonstrated that, like TMEM63B^20^, the PLS activity of TMC1 and TMC2 is regulated by cholesterol.

Cholesterol influences the mechanical properties of membranes by reducing bilayer fluidity, increasing lipid packing and rigidity, and decreasing permeability^59,60^. Therefore, it is possible that rather than acting as an agonist, cholesterol may indirectly regulate TMC lipid scrambling by altering the biophysical properties of the membrane. To get some insight on how cholesterol regulates TMC scramblase activity, we calculated the number of cholesterol molecules that CmTMC1 and MuTMC2 require to reach maximum scramblase activity based on the number of proteins per liposome we have in our system (see Methods). We estimated that TMCs require more than 300 cholesterol molecules per TMC dimer, sufficient to coat the entire lipid-exposed TMC surface, suggesting that cholesterol influences TMC lipid scrambling indirectly, likely through changes in the biophysical properties of the membrane. Consistent with this, we found that the MET blocker benzamil activated PLS activity *ex vivo* without altering the cholesterol levels in the cochlea (untreated: 4.67 ± 0.23 µg/mL and treated: 4.76 ± 0.05 µg/mL, n = 4). Additionally, Ca^2+^ has been shown to regulate membrane diffusivity^61^ and tip-link stiffness and tension^56,62^ in mammalian auditory hair cells. Therefore, it will be important to consider in future investigations the possibility that MET channel blockade, tip-links disruption, or reductions in intracellular Ca^2+^ may trigger TMC-dependent PS externalization by influencing membrane properties.

Cholesterol is the second most abundant lipid in hair cells. The plasma membrane of embryonic chick vestibular hair cells contains 43% PC, 15% PE, 6% PS, 3% sphingomyelin, 3% phosphatidylinositol phosphates, and 28% cholesterol^36^. Our data show that at these physiological levels of cholesterol, wild-type TMC proteins are unable to scramble lipids and that only negligible levels of PS externalization are detectable in healthy wild-type neonatal hair cells. However, treatment of mouse auditory hair cell explants with mβCD, which resulted in a reduction of ∼60% of cholesterol, triggered PS externalization. We observed higher levels of externalized PS in outer hair cells (OHC) compared to inner hair cells (IHC). This might be explained by the higher cholesterol content of IHC^63^ or differences in their membrane viscosity^64^. We propose that PLS is an intrinsic activity of TMC proteins that remains inactivated in healthy conditions with physiological MET operation and membrane homeostasis. In situations where the TMC MET channel function or hair cell membrane properties are compromised, the scramblase activity of TMCs is activated, PS is externalized, and membrane blebs are released, facilitating the removal of non-functional MET complexes and the renewal of stereocilia membrane (Figure 7B).

Importantly, enhancement or dysregulation of this novel TMC scramblase function can lead to pathogenesis. TMC1 (OMIM: 606706) is a major contributor to autosomal recessive nonsyndromic hearing loss, with over 125 pathogenic variants identified to date, and just eight of those linked to autosomal dominant hearing loss^65,66^. In this study, we show that two dominant (MmTMC1/CmTMC1: M412K/M507K and D569N/D658N) and one recessive (D528N/D617N) deafness-causing mutations enable TMC1 to scramble lipids in the presence of physiological cholesterol levels, resulting in constitutive externalization of PS (Figure 7B). Interestingly, the D617N mutation displays an intermediate level of PLS activity between wild-type TMC1 and the two dominant mutations, potentially explaining its recessive nature and providing a mechanistic link to the associated deafness phenotype. Mutations in similar or equivalent pore-lining residues in TMEM16s^67–69^ and TMEM63B^20^, have been shown to alter PLS activity, which suggests that the residues altered in these hearing loss-associated mutations play a direct role in TMC1 lipid scrambling. However, these mutations also change the properties of the TMC1-dependent MET channels^12–15^. Therefore, to fully understand how these mutations impact hearing and elucidate the relationship between proper TMC function and hair cell physiology, it will be important to investigate the interplay between TMC channel and scramblase function.

Mice carrying either one (dominant) or two (recessive) copies of these TMC1 mutations exhibit constitutive PS externalization, followed by hair cell loss and hearing loss, suggesting a potential link between TMC1, the maintenance of plasma membrane asymmetry and hair cell survival. Notably, PS externalization is a hallmark of apoptotic cells^22,70^, and prolonged disruption of the plasma membrane asymmetry could contribute to hair cell death. We show that cholesterol removal by mβCD treatment in hair cell explants from *Tmc1*^M412K/M412K^ or *Tmc1*^D569N/D569N^ mutant mice reduces PS externalization. This provides a proof of principle for cholesterol reduction as a potential therapeutic intervention to mitigate hair cell loss. Consistent with this, cholesterol-lowering drugs have been reported to protect hearing in patients exposed to ototoxic compounds^71,72^. Furthermore, abnormally high or low cholesterol levels have been shown to be detrimental to hearing^73–75^. Thus, our work not only identifies a novel family of lipid scramblases but also highlights the need for future studies on how lipid metabolism and dietary fats influence auditory and vestibular physiology. Understanding the TMC role in membrane regulation may help uncover molecular mechanisms underlying other TMC-related disorders, such as cancer^76^, infertility^77^ or epidermodysplasia verruciformis^78,79^.

## Materials and methods

### Plasmids

Plasmids for baculovirus expression of His-GFP-CmTMC1 (230-828) and His-GFP-MuTMC2Δ (210-775) were a gift from Motoyuki Hattori^8^. Plasmid containing His-GFP-CmTMC1 FL (1-955), His-GFP-MuTMC2 FL (1-928), His-GFP-CmTMC1 (230-828), His-GFP-MuTMC2 (211-807), M507K His-GFP-CmTMC1 (230-828), D617N His-GFP-CmTMC1 (230-828), D658N His-GFP-CmTMC1 (230-828), His-GFP-mTMC1 (135-757), and His-GFP-mTMC2 (188-783), cloned in the BamHI/NotI sites of the pFastBac1 vector were codon-optimized and synthetized by Genscript. ANO10 (TMEM16KA-c001) plasmid was a gift from Nicola Burgess-Brown (Addgene plasmid # 110267). MmTMEM16A-GFP and MMTMEM16F-GFP plasmids were a gift from Criss Hartzell^69^. Bacmids were generated by transforming DH10Bac (ThermoFisher) cells with the pFastBac1 constructs. *Spodoptera frugiperda (*Sf9) cells were then transfected using Cellfectin to produce baculovirus particles, which were harvested and stored with 2% fetal bovine serum (FBS).

### Protein expression and purification

*Sf9* cells infected with P2 baculovirus were incubated at 27°C for 1 day and moved to 20°C until >90% of cells displayed GFP signal, signifying infection. Cells were harvested by centrifugation at 3700 rpm for 10 min at 4°C, flash-frozen in liquid nitrogen, and stored at −80°C until needed. Cell pellets were resuspended in 25 mL of solubilization buffer (50 mM Tris-HCl pH 8; 150 mM NaCl; 20 mM imidazole; 1 mM ϕ3-mercaptoethanol; 1 Roche cOmplete™, Mini, EDTA-free Protease Inhibitor Cocktail tablet) without detergent, homogenized, and lysed by probe sonicator for 24 min. A 10:2 mixture of n-dodecyl-ß-D-maltoside and cholesteryl hemisuccinate (DDM/CHS) was added to the lysate for a final concentration of 2/0.4% (w/v), and rotated overnight at 4°C on a tube rotator. The next day, insoluble material was removed by 1h ultracentrifugation at 40000 rpm and 4°C, and the supernatant was incubated with 2-5 mL Qiagen Ni-NTA beads on a rotator at 4°C for at least 3 h. Beads were washed with buffer A (20 mM Tris-HCl pH 8, 300 mM NaCl, 0.05/0.01% DDM/CHS, 1 mM BME, 20 mM imidazole), and then the protein was eluted with buffer B (20 mM Tris-HCl pH 8, 300 mM NaCl, 0.05/0.01% DDM/CHS, 1 mM BME, 500 mM imidazole) in 1 mL aliquots. Eluates were further purified by size exclusion chromatography on a Superose 6 increase 10/300 GL column in buffer C (20 mM Tris-HCl pH 8, 300 mM NaCl, 0.05/0.01% DDM/CHS, 2 mM DTT) and stored at –80°C until used. Protein tolerance to freeze-thaw was verified by comparing fluorescence-detection size exclusion (FSEC) chromatograms. For purifications of TMC proteins in FC12, overnight solubilization was conducted with 0.525% FC12 (w/v). Buffers A and B were prepared with 0.1575% FC12, and buffer C was prepared with 0.07% FC12.

To obtain untagged proteins, elution fractions from the nickel column purification were incubated overnight at 4°C with Thrombin (1 unit protease per ∼20 µg TMC1 constructs) or TEV (1 unit protease per 1 µg TMC2 constructs) proteases, followed by a 2-h incubation at 25°C. Cleaved and uncleaved proteins were further separated through nickel column purification. The histidine or GFP tags did not affect the scramblase function of the proteins (Supplemental Figure 5), and functional experiments were performed with tagged proteins unless indicated.

### Fluorescence-based in vitro scramblase assays

All lipids were purchased from Avanti and dissolved in chloroform at a final concentration of 25 mg/mL. Most of our experiments were performed using liposomes containing: 1-palmitoyl-2-oleoyl-sn-glycero-3-phosphocholine (POPC, Cat 850457C), 1-palmitoyl-2-oleoyl-sn-glycero-3-phosphoethanolamine (POPE, Cat 850757C), and 1-palmitoyl-2-oleoyl-sn-glycero-3-phospho-L-serine (POPS, Cat 840034C) at a 2:1:1 ratio with 10% cholesterol (CHO, Cat 700000P) and 0.4% of the fluorescent 1-myristoyl-2-sn-glycero-3-phosphoethanolamine (NBD-PE, Cat 810151P), which mimics the stereocilia lipid content^36^. The cholesterol percentage is consistently indicated in weight/volume.

To prepare the liposomes, lipids were dried under a gentle stream of argon gas (Roberts Oxygen), washed with pentane, dried again, and resuspended to a final concentration of 20 mg/mL in reconstitution buffer (50 mM HEPES pH 7.4, 300 mM KCl) with 35 mM 3-[(3-cholamidopropyl) dimethylammonio]-1-propanesulfonate (CHAPS). After a 20-min incubation at room temperature (RT), lipid mixtures were probe sonicated (Pulse 150 homogenizer) for 5-10 min until clear and split into 200 μL aliquots. For every experiment, at least one aliquot was used as a protein-free control (liposomes). Detergent-solubilized protein (60 μg, unless otherwise indicated) was buffer-exchanged using a 50 kDa concentrator to remove DTT, resulting in purified protein in 20 mM HEPES pH 7.4, 150 mM NaCl, 0.05/0.01% DDM/CHS (or 0.063% FC-12), and 2 mM EGTA, or neither. After buffer exchange, the protein was incubated with the liposomes for 30 min at room temperature and detergent was removed using five 2h incubations with 200 mg/mL of BioBeads SM-2 (Bio-Rad) with rotation at 4°C. The third or fourth bead incubation was conducted overnight. After the final bead wash, liposomes and proteo-liposomes were subjected to sonication, freeze/thaw cycles, then extrusion through a 400 nm membrane. Fluorescence measurements were performed in a Fluoromax (HORIBA) using 10 μL of sample in 2 mL EGTA assay buffer (50 mM HEPES pH 7.4, 300 mM KCl, 2 mM EGTA). The fluorescence signal was recorded at 25°C, 4 Hz, using an Ex/Em 470/530 and slit #3 with constant gentle mixing. BSA- and SDT-PLS assays were performed in parallel using the same samples. In all PLS experiments, outliers were defined as more than 3 standard errors away from the mean, and discarded from analyses of significance.

For the heat-inactivated experiments, liposomes were prepared as described above. Following the 30-minute incubation of the protein with the liposomes or after the fifth BioBead wash samples were split in two; control (RT) and heat-inactivated (HI). Control sample was maintained at room temperature for 1h while heat inactivation was performed by incubating the sample for one hour at 95 °C in a thermoblock (Eppendorf Thermomixer C).

For the MET Blocker experiments, a final concentration of 0.1 mM MET blockers (streptomycin, neomycin, curare, or benzamil) was added to liposomes following sonication. No blockers were added to the protein-free control. Additionally, one of the proteo-liposome samples served as the negative control, with no MET blocker. The BioBead incubation and fluorescence measurements proceeded as normal.

For the Ca^2+^ experiments, dried lipids were resuspended in Ca^2+^ assay buffer (50 mM HEPES, 300 mM KCl, 0.5 mM Ca(NO_3_)_2_, pH 7.4), EGTA assay buffer (50 mM HEPES, 300 mM KCl, 2 mM EGTA, pH 7.4), or reconstitution buffer (50 mM HEPES, 300 mM KCl, pH 7.4). Sonication, addition of protein, BioBead incubation, and extrusion were conducted in the same way as described above. For the fluorescence recordings, 10 µL of sample was added to a final volume of 2 mL of Ca^2+^ assay buffer, EGTA assay buffer, or reconstitution buffer. For the Ca^2+^addition experiments, liposomes were prepared as described above, but in reconstitution buffer (without EGTA). During the fluorescence measurement, SDT was added at 100 s, and Ca(NO3)_2_ was added to a final concentration of 0.5 mM at 500 s. To assess the role of Ca^2+^at different time points, liposomes were prepared as before, and extruded samples were diluted, 10 µL in 2 mL reconstitution buffer, and Ca(NO3)_2_ was added to a final concentration of 0.5 mM. The samples were incubated for specific time points (5, 15, or 30 min) with constant stirring at 25 °C. After the incubation, the fluorescence recording was started and SDT was added at 100 s.

### Cholesterol addition to liposomes

For cholesterol addition experiments, POPC liposomes and proteo-liposomes were prepared as described above, except with 18:1-12:0 NBD-PE (Avanti 819156) rather than 14:0-06:0 NBD-PE to reduce NBD lipid extraction by methyl-β-cyclodextrin (mβCD). After the final extrusion, the scramblase activity of the samples was recorded before proceeding with cholesterol treatment.

Complex of mβCD with cholesterol (mβCD/CHO)^44^ were prepared concurrently with the liposomes and proteo-liposomes for each experiment. Four mβCD/CHO solutions (1, 2, 5 and 10) containing 5 mM mβCD and increasing amounts of cholesterol were prepared by transferring 100, 250, 500, or 1000 µL of a 10 mM solution of cholesterol (Sigma C8667) in chloroform to glass tubes. After drying under an argon stream, 5 mL of a 5 mM (6.55 mg/mL) solution of mβCD (Sigma 332615) in scrambling reconstitution buffer was added to each sample, followed by sonication in a bath sonicator (37 Hz, 100% power, pulse mode) for 15 min. Samples were then incubated in a shaker at 225 rpm, overnight at 35°C, after which they were sonicated for an additional 5 min, and filtered through a 0.22 µm filter.

To increase the cholesterol content, 50-80 µL of liposome or proteo-liposome samples lacking cholesterol were incubated with 500-800 µL of mβCD (control) or the four mβCD/CHO solutions (1:10 v/v for liposome or proteo-liposome samples to treatment) for 45 min at 30°C, 1000 rpm in a tabletop shaker. Vesicles were collected by ultracentrifuging samples at 50,000 rpm for 45 min in a P50A3_5049 rotor (Eppendorf). The supernatant was discarded and replaced with 50-80 µL of reconstitution buffer, and the pellet was re-dissolved using a mini-pestle. Since treatment can also extract NBD lipids, 20 µL of sample rather than 10 µL was diluted in 2 mL of assay buffer for the fluorimeter recordings. Scramblase activity is always reported as the difference between the fluorescence quenching in proteo-liposome samples compared to the corresponding liposome samples.

### Analysis of fluorimeter traces

An in-house python script was used to analyze all fluorescence traces for scrambling. Scrambling curves were fit to a function (1) that describes a flat baseline (*F*_max_) that begins to decay mono-exponentially to a second baseline (*F*_plateau_) following SDT addition at *t* = *t*_SDT_.

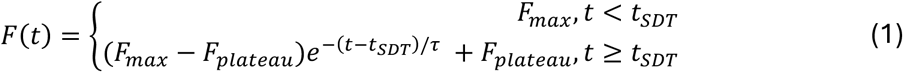

Then, the data was normalized by the fitted F_max_, and fit to the following function (2) to re-obtain the SDT addition time (*t*_SDT_):

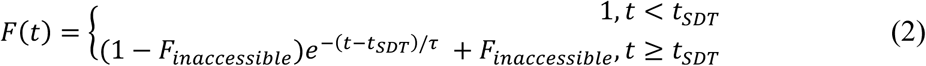

A final fit (3) of the data after *t*_SDT_ yielded the fraction of NBD lipids inaccessible to SDT (*F*_inaccessible_) as well as the time constant (τ) of the process.

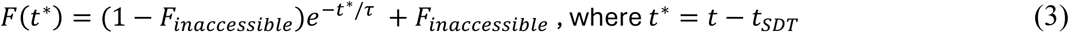

The same process was used to fit BSA data. The reported scramblase activity was calculated (4) as:

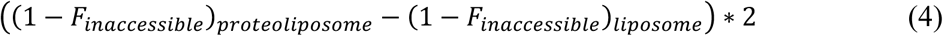

The factor of 2 scales the scramblase activity such that the maximal possible activity is 1, or 100%.

Analysis of the kinetics of NBD fluorescence loss after SDT addition revealed that the process was best modeled by a single-exponential decay with a time constant (τ) of ∼5-10 s in our standard experimental conditions, consistent with previous reports^33,80^. This rate of decay was comparable in liposomes and TMC proteo-liposomes (6.38 ± 0.5 s for CmTMC1 and 8.79 ± 0.91 s for MuTMC2) and within the range of the expected time constant for SDT quenching of NBD lipids; thus, we could not fit a lipid scrambling rate, and can only conclude that the TMC scrambling rate is faster than the NBD reduction rate by SDT. This would suggest that each functional unit of TMC can scramble >10^4^ lipids/s, which would speed up the intrinsically slow translocation of phospholipids across the membrane leaflets from more than 10 h to less than 10 s.

### Cholesterol content quantification

Cholesterol was quantified using the Amplex Red Cholesterol Assay Kit (Invitrogen A12216), which provides a fluorometric measure of cholesterol and cholesterol ester content. A standard curve was generated for every experiment, and samples were diluted in the provided Amplex assay buffer to ensure they were within the linear component of the fit (< 10 μg/mL cholesterol). The lower limit of this assay is 200 nM (80 ng/mL) cholesterol.

Samples used for cholesterol experiments were quantified for their cholesterol content. The scramblase activity and the cholesterol content were averaged across each condition, categorized by protein, purifying detergent, lipid composition, and target cholesterol content (0%, 1%, 2%, 5%, and 10%) or mϕ3CD/CHO treatment (control, 1%, 2%, 5%, and10%). These averages for each protein were then fit to an exponential plateau function (5). The cholesterol content at which the proteins reach half of the maximal activity is reported. Note that the activity at 0 cholesterol is not necessarily 0.

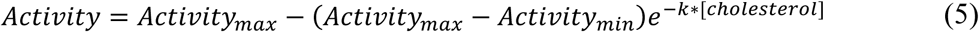

To extract lipids from cochlear explants, 2-4 cochleae were dissected from P6 mice and treated with 200 µL of a mixture of chloroform:isopropanol:NP-40 (7:11:0.1) and dissolved with a mini-homogenizer. The samples were then centrifuged (15,000 g for 10 min at 4°C), and the supernatant was discarded. The pellet was dried on a speed vac (Eppendorf) for at least 10 min to remove trace amounts of organic solvent and stored at −80C until cholesterol quantification. To quantify the cholesterol in these samples, the dried lipids were resuspended in 50 µL of the Amplex assay buffer and sonicated in the water sonicator for 5 min before proceeding with the Amplex treatment as explained above. The cholesterol content of organs of Corti was measured for samples from wild-type and *Tmc1*^M412K/M412K^ mutant mice under several conditions.

### Liposome size estimates

Dynamic light scattering (DLS) analysis was conducted to estimate the size of the liposomes used for scramblase assays. Fresh scrambling samples were diluted 1:500 in EGTA assay buffer for DLS measurements. Each measurement represents the average of 10-20 recordings on a Dynapro Nanostar instrument. The autocorrelation curve was subjected to a cumulant fit to obtain the particle size and the polydispersity. The analysis revealed the presence of homogenous liposome and proteo-liposome populations with a radius of 96.7 ± 5.2 nm for protein-free liposomes, 104.4 ± 6.6 nm for CmTMC1 proteo-liposomes, and 109.4 ± 4.0 nm for MuTMC2 proteo-liposomes. Assuming that the average molecular weight of the reconstituted proteins is 99.2 kDa for TMC1 and 98.5 kDa for TMC2, that the average diameter of the reconstituted vesicles is 100 nm, and that reconstitution follows a uniform distribution, each vesicle would contain around 11-12 TMC proteins on average^33^.

### All-atoms MD simulations

Dimeric structural models for truncated CmTMC1 and MuTMC2 were generated using AlphaFold3^46^ and embedded into mixed membrane lipid bilayers of POPC:POPE:POPS at a 2:1:1 ratio with or without 10% cholesterol using CHARMM-GUI^81,82^, as summarized in Supplemental Table1. The protein-lipid systems were solvated using TIP3P explicit water and neutralized with 0.3 M KCl.

All-atom equilibrium MD simulations were performed using NAMD 3.0^49^ and the CHARMM36 force field with CMAP corrections^81,83^. For all systems, equilibrium simulations consisted of 2,000 steps of minimization, 0.5 ns of dynamics with all components except lipid tails fixed or constrained, 0.5 ns of dynamics with harmonic constraints applied to the protein (1 kcal mol^-1^ Å^-2^), 1 ns of unrestrained dynamics in the *NpT* ensemble (*γ* = 1 ps^-1^), and up to 100 ns of free dynamics in the *NpT* ensemble (*γ* = 0.1 ps^-1^) prior to non-equilibrium production runs. A piston period of 200 fs and a damping timescale of 50 fs was used for pressure control and atomic coordinates were saved every 5 ps. All simulations were performed at 310 K and 1 atmosphere of pressure by using the Langevin thermostat and Nosé-Hoover Langevin pressure control. A 12 Å cutoff distance was employed with a force-based switching function starting at 10 Å. Periodic boundary conditions and the PME method were used to calculate long-range electrostatic interactions with a grid density greater than 1 Å^-3^. An integration timestep of 2 fs with electrostatic interactions computed every other timestep was utilized for every simulation in tandem with the SHAKE algorithm to constrain hydrogen atoms.

Following equilibrium MD simulations, we did TMD simulations on all four systems for 20 ns to induce pore opening in TMC models^49,50^. The simulations were guided by a model of the MmTMC1 protein previously generated based on open state structures of TMEM16 proteins^18^. To match each simulated TMC variant, reference structures for TMD simulations were generated by mutating the template residues in COOT. For the CmTMC1 systems, residues selected as TMD targets included aligned regions within transmembrane helices TM3 (residues 222–258; numbering according to MmTMC1), TM4 (268–293), TM5 (298–327), TM6 (375–410), TM7 (432–451), and TM8 (455–477). For the CmTMC2 systems, corresponding targeted residues encompassed aligned positions in TM3 (219–258), TM4 (268–289), TM5 (294–323), TM6 (372–408), TM7 (430– 452), and TM8 (455–475). In these simulations, Cα-atoms root mean square deviation from the target model was minimized from ∼ 6 Å to 0 Å. In all four systems, one subunit of each dimeric TMC model was maintained in the closed-pore conformation predicted by AlphaFold3, as a control. Following TMD simulations, all four systems listed in Supplemental Table1 underwent additional equilibrium MD simulations up to 1.7 μs, during which harmonic positional constraints (*k* = 2 kcal mol⁻¹ Å^-2^) were applied to transmembrane helices TM3, TM4, TM5, TM6, TM7, and TM8 of one subunit to keep its pore open. Unless otherwise specified, all remaining simulation parameters were as previously described.

### CG MD simulations

Following completion of all-atom minimization, equilibration, and TMD simulations, atomistic solvent (water and ions) and lipid molecules were removed, and the protein models were converted to the Martini 3 CG representation using Martinize2/Vermouth (vermouth-0.10.1.dev93)^84^. Secondary structure information required for the Martini CG mapping was calculated with DSSP^85^ integrated within MDTraj^86^. For all CG TMC systems, one pore was kept open, elastic network bonds were implemented (*k* = 700 kJ·mol^-1^·nm^-2^; distance bounds: 5–9 Å), and elastic bonds between monomeric subunits were not removed. The CG systems were constructed using Insane (version 1.2.0)^87^, incorporating a lipid bilayer composed of POPC, POPE, and POPS at a 2:1:1 ratio, supplemented with an additional 10% cholesterol when applicable. Regular-sized water beads (W) were added to achieve box dimensions of approximately 180 × 180 × 220 Å³. Monovalent ion beads (Na^+^ and Cl^-^) were included to neutralize the system and achieve a final ionic strength of 0.3 M. The vertical (*z*-axis) placement of the protein within the membrane was adjusted according to predictions from the positioning of proteins in membrane (PPM) 3.0 server^88^.

CG MD simulations were conducted using GROMACS version 2024.2^89^ with the Martini 3 *New-Reaction-Force (RF)* parameter set^90^. All simulations employed a 20-fs integration timestep and utilized RF electrostatics. Pressure was regulated at 1 bar semi-isotropically (compressibility: 3 × 10⁻⁴ bar⁻¹) using the Berendsen barostat^91^ (coupling constant: 4 ps⁻¹), while the temperature was maintained at 310 K using the velocity-rescale (v-rescale) thermostat^92^. To prevent simulation artifacts associated with infrequent neighbor list updates^93^, Verlet-buffer-tolerance was disabled, a conservative neighbor list cutoff of 1.5 nm was adopted, and neighbor searching (nstX variables) was set to occur every 20 simulation steps, in accordance with recommendations for Martini 3 *New-RF*. System equilibration followed the following steps: (1) Up to 500 steps of position-restrained energy minimization in vacuum, (2) Up to 500 steps of position-restrained minimization after solvating the system with water, ions, and lipids, (3) 10 ps of position-restrained *NVT* equilibration with a 2-fs timestep, and (4) subsequent *NpT* equilibration phases without positional restraints, lasting 25 ps (5 fs timestep), 50 ps (10 fs timestep), and 300 ps (20 fs timestep), respectively. Following equilibration, production simulations were extended to 10 µs (Supplemental Table1), with atomic positions recorded every 100 ps for analysis.

Scrambling events in all-atom and CG MD systems were identified and monitored through visual inspection. Trajectory analyses, molecular visualization, and video generation for both all-atom and CG systems were performed using VMD^94^, and data plotting was carried out using Xmgrace and Origin software. Additionally, positional data along the membrane normal (*z*-direction) were extracted using VMD to track protein movements during scrambling events.

### Cell-based scramblase assays

HEK293 cells (ATCC) growing in Dulbecco’s Modified Eagle Medium (DMEM) without Ca^2+^ supplemented with 10% fetal bovine serum (FBS) and GlutamaX, were seeded at 70% confluency in collagen-coated 12-well plates. The next day, cells were transfected with FUGENE (Promega) and the corresponding cDNA to transiently express GFP-tagged MmTMEM16A, MmTMEM16F, or MmTMC1. Approximately 16 h post-transfection, cells were washed with fresh media and recovered for 4 h at 37 °C in the cell incubator. Cell media was removed and the Ca^2+^ ionophore A23187 was added to a final concentration of 10 μM for 2-5 min. Media was removed and cells were washed once with buffer 1 (10 mM Tris HCl pH 7.4 and 130 mM NaCl) and with buffer 2 (10 mM Tris HCl pH 7.4, 130 mM NaCl, and 2.5 mM Ca_2_Cl). Annexin V diluted at 1:20 in buffer 2 was added to the cells and incubated for 10 min at RT. Cells were then washed twice with buffer 2 and fixed in 2% PFA in buffer 2 for 20 min at RT. Coverslips containing cells were mounted on an imaging slide and imaged in a confocal microscope.

### Hair cell experiments

All animal experiments were performed in accordance with the NIH guidelines and were approved by the NIH Office of Animal Care and Use NIDCD (animal protocol number #1617). Wild-type C57BL/6J mice were obtained from The Jackson Laboratory (strain #000664), while *Tmc1*^D569N/D569N^ were a gift from Robert Fettiplace and Tmc*1*^M412K/M412K^ from Wade Chien and Andrew Griffith.

Temporal bones were isolated from 6-days old wild-type C57BL/6J, *Tmc1*^D569N/D569N^ or *Tmc1*^M412K/M412K^ mice^12,17^ and dissected to expose the organ of Corti in cold Leibovitz’s L15 medium. Auditory hair cells explants were transferred into a Corning PYREX 9-well depression plate and incubated for 30 min at RT under gentle orbital shaking and protected from light in HBSS (control), HBSS containing 0.1 mM benzamil (benzamil), 10 mM mβCD in HBSS (mβCD), or HBSS containing 0.1 mM benzamil and 10 mM mβCD (mβCD + benzamil). 1:25 annexin V–Alexa Fluor 647 (Thermo Fisher Scientific) was included in all these conditions to label and visualize externalized PS. After treatment, tissues were washed twice with HBSS and fixed in 4% paraformaldehyde (PFA) in HBSS for 30 min and explants were stained for 1 h with 1:1000 spy555-actin (Cytoskeleton, Inc.) in PBS for actin filaments visualization. The spiral ligament and tectorial membrane were carefully removed after fixation, leaving only the stained organ of Corti explants in HBSS buffer. Tissues were washed with phosphate-buffered saline (PBS) to remove excess salt and subsequently mounted in ProLong Diamond antifade mounting medium (Thermo Fisher Scientific) on Superfrost Plus microscope slides (Fisherbrand). Coverslips (No. 1.5, 0.17 ± 0.02 mm thickness, Warner Instruments) were applied for confocal imaging.

Imaging was performed using a confocal laser scanning microscope (Zeiss LSM 980, Carl Zeiss AG) equipped with a 32-channel Airyscan2 detector, an oil immersion alpha Plan-Apochromat 63×/1.4 Oil Corr M27 objective (Carl Zeiss), and immersol 518F medium [ne = 1.518 (30°)]. Control and treated samples from each experiment were imaged on the same day, using identical image acquisition settings, with no averaging, and optimized parameters for *x*, *y*, and *z* resolution for each independent experiment. Image acquisition was conducted with Zen Black 2.3 SP1 software (Carl Zeiss), and microscopy data analysis and quantification were performed in ImageJ. To measure annexin V fluorescence intensity at the apical region of the hair cells, a region of interest (ROI) was defined around each OHCs and IHCs using the elliptical tool as represented in Figure 4E. The fluorescence intensity of an equivalent ROI in an area outside the hair cells was considered background and subtracted from the values measured in the hair cells for each image. Graphs and statistical analysis were done in GraphPad Prism unless otherwise indicated.

## Acknowledgments

We would like to thank Alessio Accardi and Zhang Feng for their assistance with the *in vitro* scramblase assays, and Wei-Hsiang Weng for valuable suggestions on TMD simulations; Kenton Swartz, Katherine Kindt, Sergio M. Pontejo, and Catherine Weisz for their valuable comments on the manuscript; and Criss Hartzell, Motoyuki Hattori, Robert Fettiplace, Andrew Griffith and Wade Chien for sharing cDNAs and mouse strains with us. We also thank the NHLBI Biophysics Core for providing assistance with the DLS experiments, and the NINDS Mass Spectrometry Core for sample analysis. Simulations were performed on the Polaris supercomputer at the Argonne Leadership Computing Facility (2025 DOE/ALCF INCITE project: MechanoEar) and on Stampede 3 (NSF/ACCESS grant MCB140226). This work was supported by the Division of Intramural Research of the National Institute on Deafness and Other Communication Disorders (NIDCD DIR DC000096 to AB) and by the National Institutes of Health (NIH R01 DC015271 to MS).

## Author contributions

Conceptualization: AB

Data curation: YCP, AB, MS

Formal analysis: RC, YCP, AB, MS, HW Funding acquisition: AB and MS Investigation: HL, YCP, JB, RC, HW, HES

Project administration and Supervision: AB and MS Validation: HL, YCP, JB, RC, AB

Visualization: YCP, HL, AB, HW, MS Writing – original draft: AB

Writing – review & editing: YCP, HW, MS, AB

## Resource availability

All unique reagents generated in this study or additional information required to reanalyze the data reported in this paper are available from the lead contact. Data reported in this paper and original code to analyze the PLS assays are available at Dryad.

## Supplemental information

### Supplementary Movies

**Movie S1 – Lipid scrambling during an all-atom MD simulation of CmTMC1.** A scrambling event observed through CmTMC1 (red) in the presence of 10% cholesterol. A POPE (green) molecule translocated through the open pore during the ∼ 1.7-μs long equilibrium MD simulation. Other lipids head groups are shown as white spheres. Protein is shown as a red molecular surface. Water molecules, ions, and lipid tails for non-translocating lipids are omitted for visualization purposes.

**Movie S2 – Lipid scrambling during an all-atom MD simulation of MuTMC2**. Two scrambling events observed through MuTMC2 in the presence of 10% cholesterol, where both POPC (gray) and POPE (green) molecules translocated through the open pore during the 1.7-μs long equilibrium MD simulation. System shown as in Movie S1 with protein in blue.

**Movie S3 – Canonical lipid scrambling during a CG MD simulation of MuTMC2.** A POPC lipid (1207) translocated through the open pore subunit of the CG MuTMC2 dimeric complex without cholesterol. Most translocation events are through the open pore. System shown as in Movie S2.

**Movie S4 – A rare non-canonical scrambling event during a CG MD simulation of CmTMC1.** A POPC lipid (807) translocated through the closed pore subunit of the CG CmTMC1 dimeric complex without cholesterol. System shown as in Movie S1.

**Movie S5 – A rare non-canonical scrambling event during a CG MD simulation of MuTMC2.** A POPC lipid (1174) translocated through the TM10 dimerization interface of the CG MuTMC2 dimeric complex in the presence of 10% cholesterol. System shown as in Movie S2.

**Movie S6 – A rare non-canonical scrambling event during a CG MD simulation of MuTMC2.** A POPS lipid (1371) translocated through the TM10 dimerization interface of the CG MuTMC2 dimeric complex in the presence of 10% cholesterol. System shown as in Movie S2.

### Supplementary Figure

**Supplementary figure 1:**
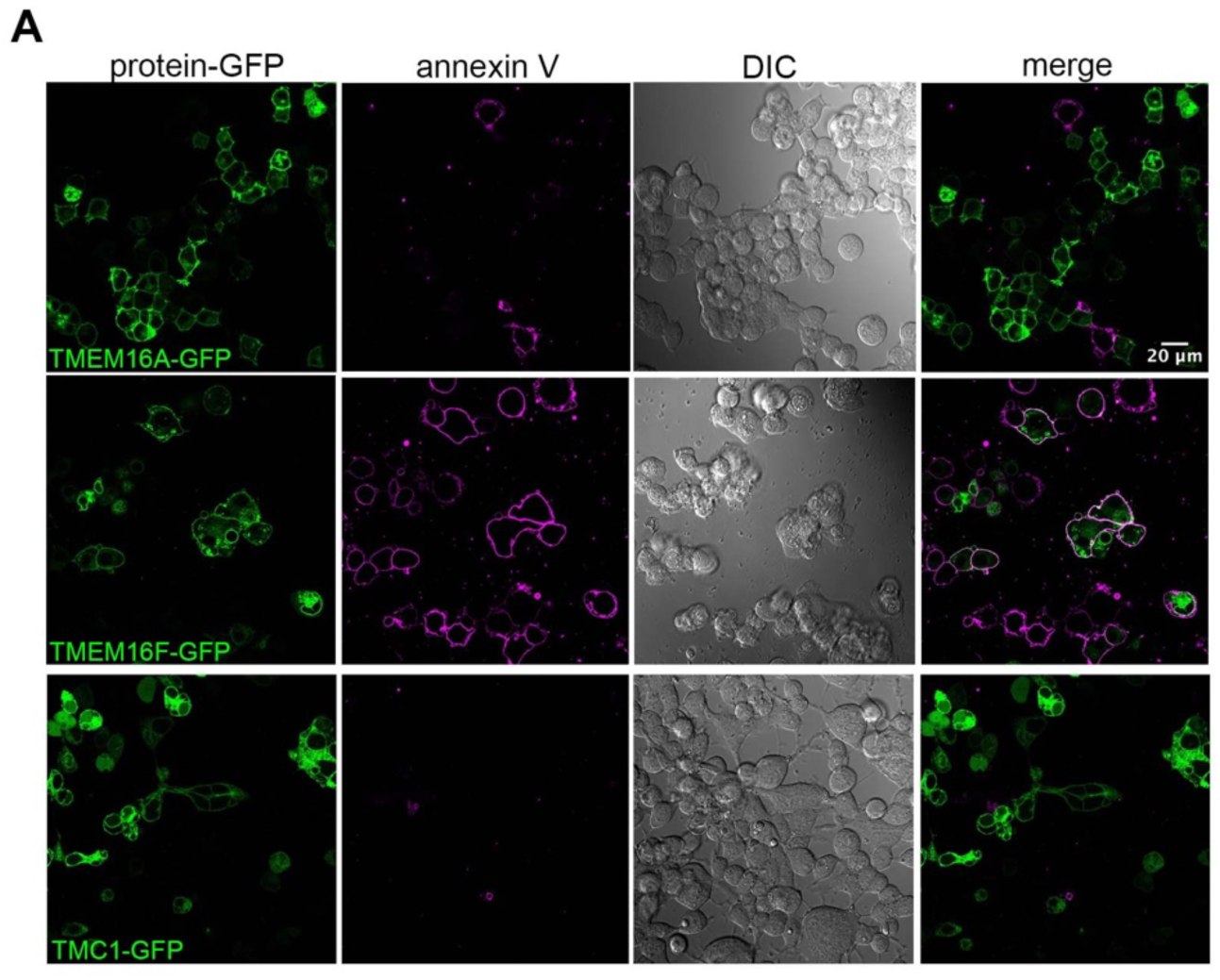
TMC1 does not trigger PS externalization when expressed in HEK293 cells. HEK293T cells transiently expressing the chloride channel TMEM16A or mTMC1 tagged with GFP (green) lacked externalized PS as observed by the absence of annexin V labeling (magenta). However, HEK293T cells expressing the lipid scramblase TMEM16F showed externalized PS at the plasma membrane that was detected by annexinV-647, indicating that TMEM16F function as a lipid scramblase. DIC is also included to appreciate the presence of non-transfected cells.

**Supplementary figure 2:**
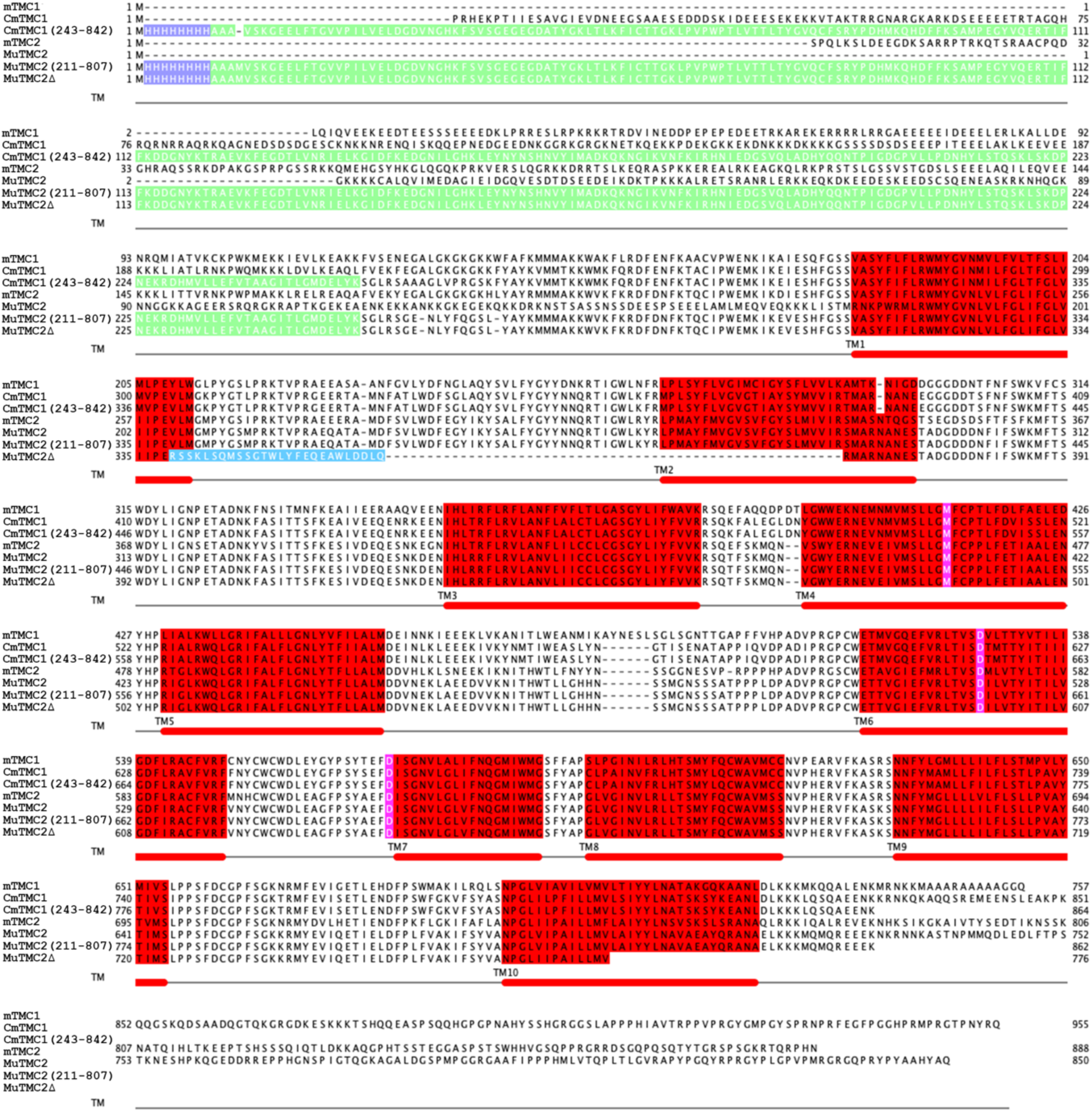
Sequence alignment of the TMC1/2 proteins used in this study. Sequence alignment of mTMC1, CmTMC1 full-length, CmTMC1 lacking the N- and C-terminals (243-842), mTMC1, MuTMC2 full-length, MuTMC2 lacking the N- and C-terminals (211-807), and MuTMC2Δ. Histidine tag is highlighted in purple, GFP in green, TM segments in red and the insertion present in MuTMC2Δ in blue. TM segments are also indicated below the alignment and deafness-causing mutations are shown in magenta.

**Supplementary figure 3:**
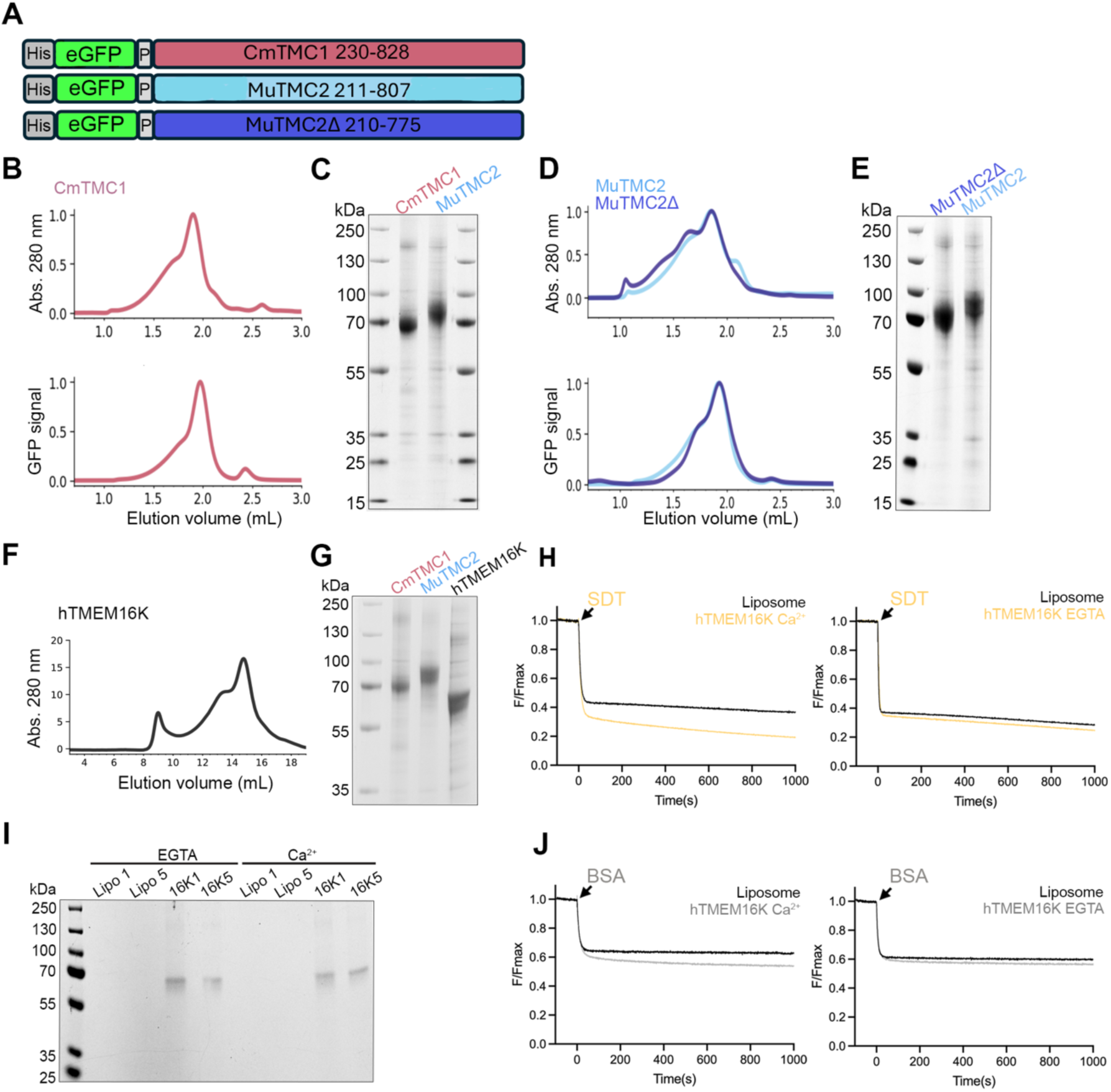
Protein purification and hTMEM16K scramblase assays. A. Cartoon representing the CmTMC1, MuTMC2 and MuTMC2Δ constructs used in this study. B. FSEC profiles showing the normalized absorbance at 280 nm (top panel) or GFP signal (bottom panel) of purified CmTMC1 in DDM/CHS. C. SDS-PAGE gel showing the CmTMC1 and MuTMC2 proteins used in our experiments. Protein bands and estimated molecular weights for the protein ladder are indicated. D. FSEC profiles showing the absorbance at 280 nm (top panel) and GFP signal (bottom panel) of purified MuTMC2 (light blue) and MuTMC21′ (dark blue) in DDM/CHS. E. SDS-PAGE gel showing the MuTMC2 (light blue) and MuTMC21′ (dark blue) proteins used in our experiments. F. FSEC profile showing the absorbance at 280 nm of purified hTMEM16K in DDM/CHS. G.SDS-PAGE gel showing purified CmTMC1, MuTMC2 and hTMEM16K used in our experiments. H-J. Fluorescence intensity decay over time due to SDT (yellow) or BSA (gray) addition to POPC/PE/PS liposomes (black trace) or hTMEM16K proteo-liposomes in the presence of 0.5 mM Ca^2+^ (left) or 2 mM EGTA (right). I. SDS-PAGE gels of representative liposome (Lipo1/5, indicating wash 1 or 5) or hTMEM16K proteo-liposome (16K1/5) samples used in our experiments, confirming the presence of protein in the proteo-liposomes

**Supplementary figure 4:**
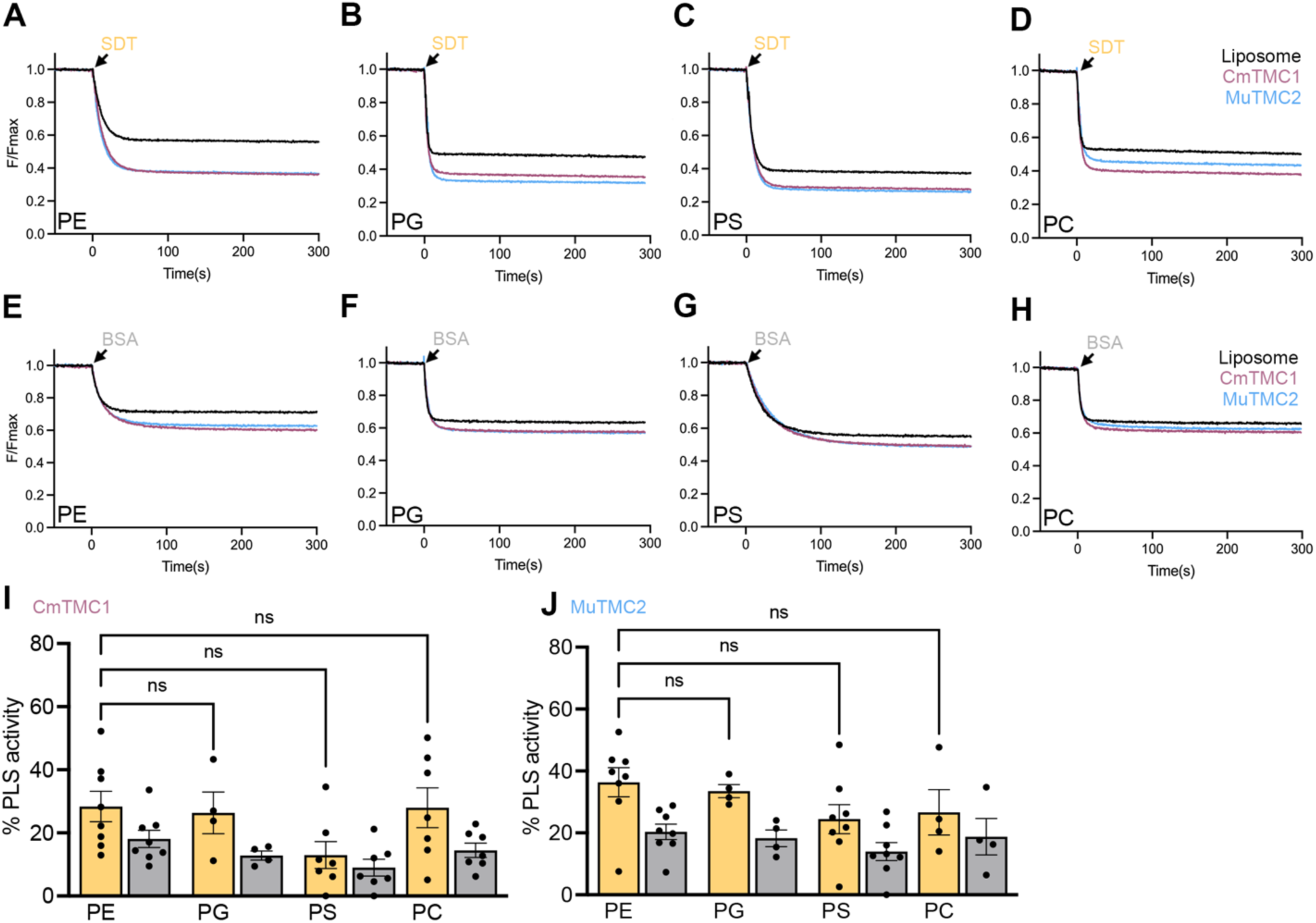
CmTMC1 and MuTMC2 scramble phospholipids with different head groups. A-D. Representative traces of NBD fluorescence decay after SDT addition to POPC/PE/PS/10CHO liposomes (black) containing 0.4% of acyl-NBD-labeled PE (A), PG (B), PS (C), or PC (D) or proteo-liposomes containing CmTMC1 (red) or MuTMC2 (blue). E-H. Representative traces of NBD fluorescence decay after BSA addition to the same samples from (A-D). I. Quantification of CmTMC1 SDT-(yellow) and BSA (grey)-based scramblase activity in the conditions shown in A-H. J. Quantification of MuTMC2 SDT-(yellow) and BSA (grey)-based scramblase activity in the conditions shown in A-H. Each dot represents one independent experiment. Mean ± SEM is shown for 3-7 independent experiments. One-way ANOVA was performed to evaluate statistical significance between NBD-PE and the other NBD-lipids. K. SDS-PAGE gels of representative liposome (Lipo1-5) or TMC proteo-liposomes (W1-5) samples used in our experiments, confirming the presence of protein in the proteo-liposomes samples.

**Supplementary figure 5:**
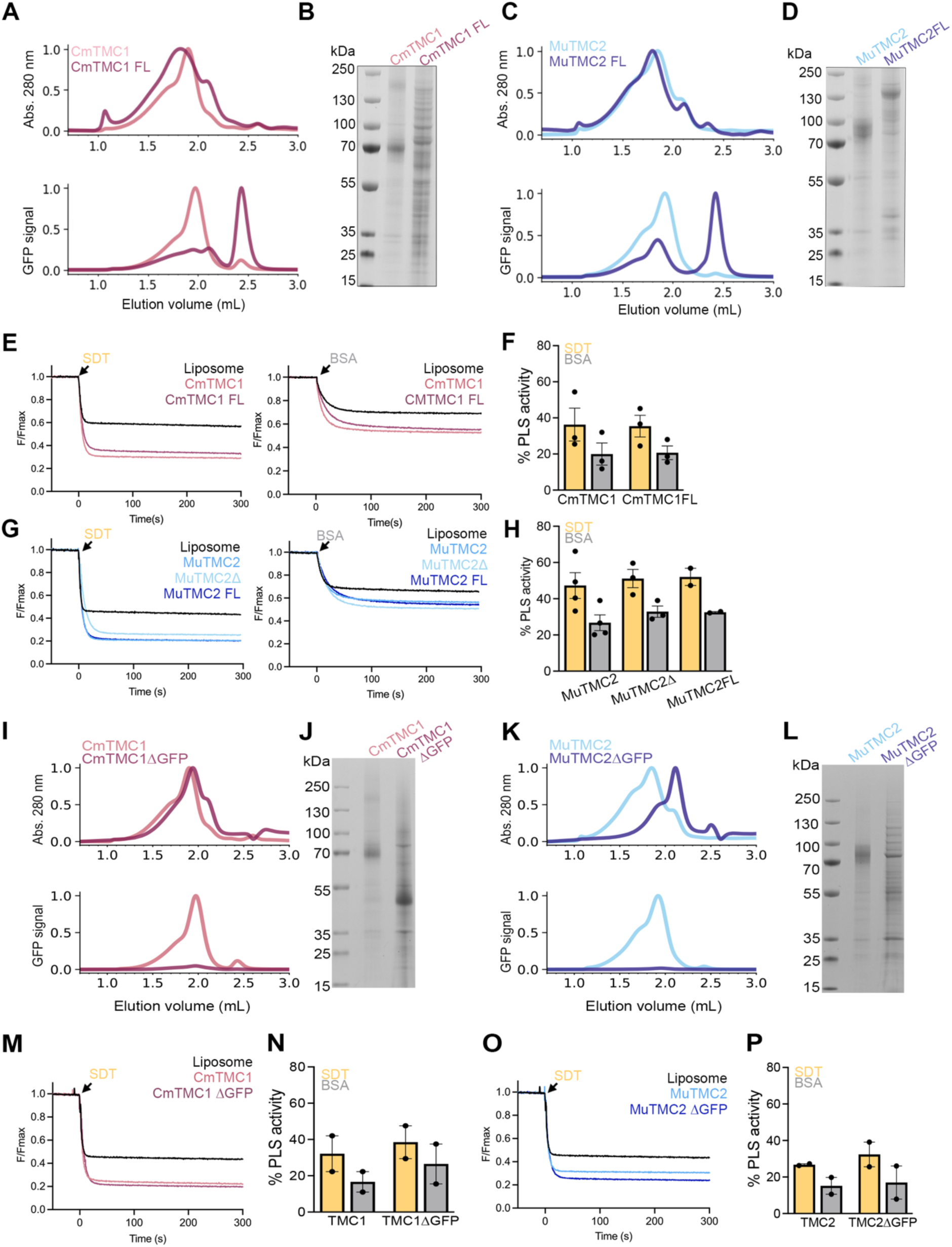
TMC scramblase activity resides at the TMC core. A. FSEC profiles showing the normalized absorbance at 280 nm (top) or GFP signal (bottom) of purified truncated (light red) or full-length (FL, dark red) CmTMC1. B. SDS-PAGE gel of the proteins shown in (A). Protein bands and estimated molecular weights for the protein ladder are indicated. C. FSEC profiles showing the normalized absorbance at 280 nm (top) or GFP signal (bottom) of purified truncated (light blue) or FL MuTMC2 (dark blue). D. SDS-PAGE gel of the proteins shown in (C). E. Fluorescence intensity decay over time due to SDT (left) or BSA (right) addition to POPC/PE/PS/10CHO liposomes (black trace) or proteo-liposomes containing truncated (light red) or full-length (dark red) CmTMC1. F. Quantification of the SDT-(yellow) or BSA-(grey) scramblase activity of truncated or FL CmTMC1. G. Fluorescence intensity decay over time due to SDT (left) or BSA (right) addition to POPC/PE/PS/10CHO liposomes (black trace) or proteo-liposomes containing truncated truncated (MuTMC2), previously published MuTMC21′, or MuTMC2 FL. H. Quantification of the SDT-(yellow) or BSA-(grey) scramblase activity of the MuTMc2 constructs shown represented in G. Mean ± SEM is shown. Each dot represents one independent experiment. One-way ANOVA was performed to evaluate statistical significance between the SDT activity of the TMC proteo-liposomes, but no statistical significance was found in F or H. I. FSEC profiles showing the normalized absorbance at 280 nm (top) and GFP signal (bottom) of CmTMC1(light red) or tagless CmTMC1 (ΔGFP, dark red). J. SDS-PAGE gel of the proteins shown in (I). K. FSEC profiles showing the normalized absorbance at 280 nm (top) and GFP signal (bottom) of MuTMC2 (light blue) or tagless MuTMC2 (ΔGFP, dark blue). L. SDS-PAGE gel of the proteins shown in (K). M. Fluorescence intensity decay over time due to SDT addition to POPC/PE/PS/10CHO liposomes (black) or proteo-liposomes containing CmTMC1 (light red) or CmTMC1 ΔGFP (dark red). N. Quantification of the scramblase activity of CmTMC1 or CmTMC1 ΔGFP. O. Fluorescence intensity decay over time due to SDT addition to POPC/PE/PS/10CHO liposomes (black) or proteo-liposomes containing MuTMC2 or MuTMC2 ΔGFP. P. Quantification of the scramblase activity of MuTMC2 or MuTMC2 ΔGFP. Mean ± SEM is shown. Each dot represents one independent experiment. One-way ANOVA was performed to evaluate statistical significance between the SDT activity of the TMC proteo-liposomes, but no statistical significance was

**Supplementary figure 6:**
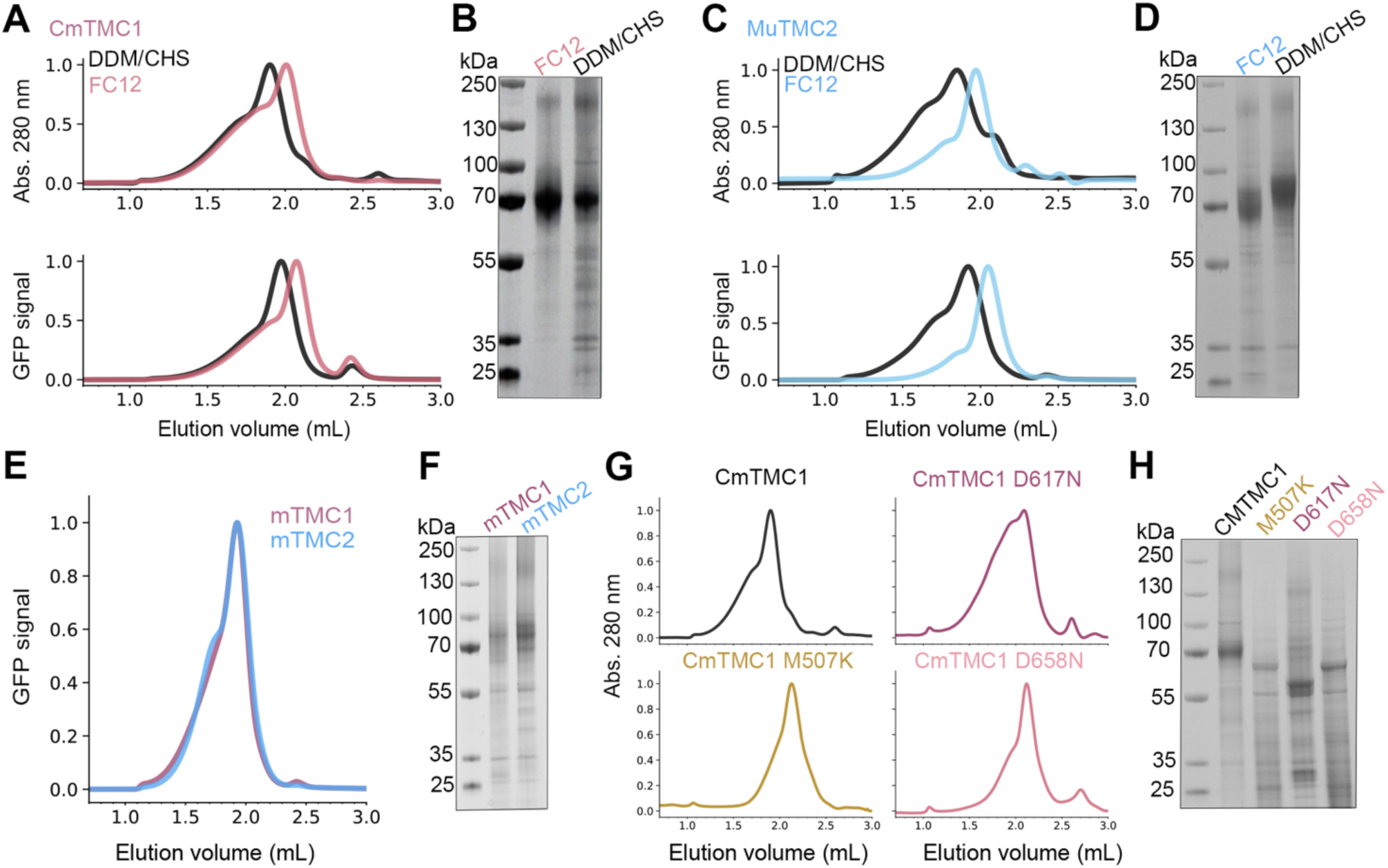
Purification of CMTMC1 and MuTMC2, mouse TMC1 and TMC2 and CmTMC1 deafness-causing mutations. A. FSEC profiles showing the normalized absorbance at 280 nm (top) and GFP signal (bottom) of purified CmTMC1 in FC12 (pink) or DDM/CHS (black). B. SDS-PAGE gel of the CmTMC1 proteins shown in (A). Protein bands and estimated molecular weights for the protein ladder are indicated. C. FSEC profiles showing the normalized absorbance at 280 nm (top) and GFP signal (bottom) of purified MuTMC2 in FC12 (blue) or DDM/CHS (black). D. SDS-PAGE gel of the MuTMC2 proteins shown in (C). E. FSEC profiles showing the normalized absorbance at 280 nm (top) of purified mTMC1 (pink) and mTMC2 (blue) used in our experiments. F. SDS-PAGE gel of the proteins shown in (E). G. Fluorescence intensity decay over time due to BSA addition to POPC/PE/PS liposomes (black) without (right) or with 10% cholesterol (left) or proteo-liposomes containing mTMC1 (red) or mTMC2 (blue). H. FSEC profiles showing the normalized absorbance at 280 nm (top) of purified CmTMC1 mutants used in our experiments. I. SDS-PAGE gel of the proteins shown in (H).

**Supplemental Figure 7:**
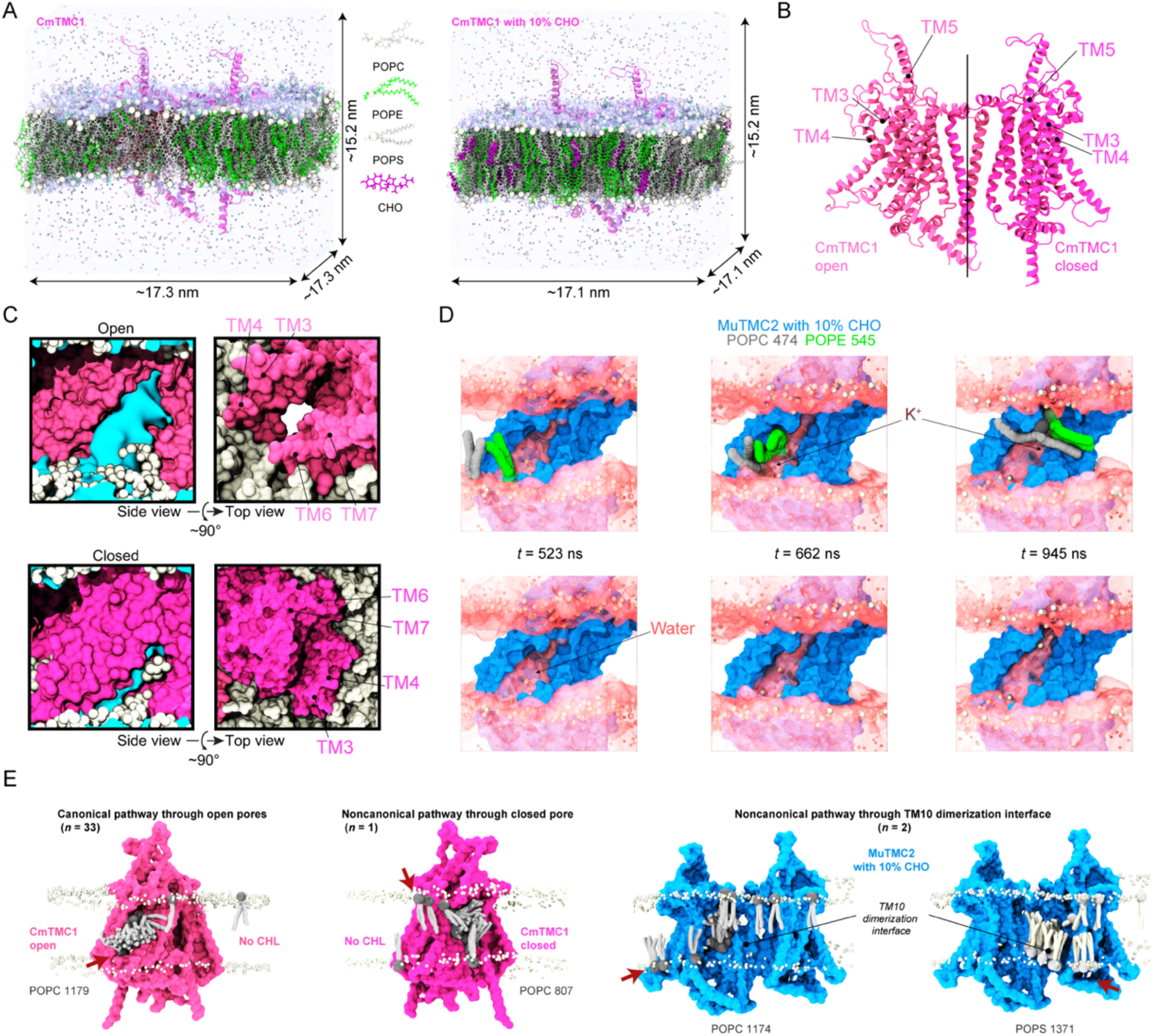
MD simulations reveal lipid scrambling by CmTMC1 and MuTMC2 through open-like groove. A. AlphaFold3-predicted dimeric model of CmTMC1 embedded into a mixed lipid POPC/PE/PS bilayers without (left) and with (right) 10% cholesterol. Similar systems were built for MuTMC2. Proteins are shown in ribbon representation, lipid phosphate atoms are shown as white spheres, water is transparent, and ions are shown as spheres. Lipids are shown as indicated in inset. B. Illustration of CmTMC1 after TMD-induced opening of one pore. C. Side and top views of one subunit of CmTMC1 where TMD simulations were used to open the pore region (top) or with a closed pore (bottom). Protein is shown in molecular surface representation, lipid headgroups are shown as white spheres, and water is in cyan. D. Snapshots at indicated time points during the lipid translocation events in the all-atom MuTMC2/10CHO system. Potassium ions (dark grey) and water molecules (red) occupied the open groove as the lipids translocated. E. Canonical (left) and rare noncanonical scrambling events (three right panels), including a scrambling event where POPC 807 translocates through the closed pore CmTMC1 and two other events (POPC 1174 and POPS 1371) showing scrambling through the TM10 dimerization interface in MuTMC2 with 10% cholesterol. In each case a translocating single lipid is shown at different time points overlaid on a representative structural model of the protein. Non-translocating lipid phosphate beads are shown as white spheres. Other system components are omitted for visualization purposes.

**Supplementary Table 1:** Summary of simulations and scrambling events.

| Label | System | $t_{\text{sim}}(\text{ns})$ | Type | Start | Size<br>(#atoms/<br>beads) | Initial Size<br>(nm<sup>3</sup>) | Number of<br>Scrambling<br>events | Scrambling<br>rates<br>(lipids·s<sup>-1</sup>) |
| --- | --- | --- | --- | --- | --- | --- | --- | --- |
| S1a | CmTMC1<br>POPC /<br>POPE /<br>POPS | 100 | EQ | - | 408,861 | 17×17×15 | 0 | 0 |
| S1b |  | 20 | TMD | S1a |  |  | 0 | 0 |
| S1c |  | 1,500 | TMD-CNST | S1b |  |  | 0 | 0 |
| S1d |  | 10,000 | CG-EQ | S1b | 68,366 | 18×18×22 | 3 | 0.3×10<sup>6</sup> |
| S2a | CmTMC1<br>POPC /<br>POPE /<br>POPS /<br>10CHO | 100 | EQ | - | 403,998 | 17×17×15 | 0 | 0 |
| S2b |  | 20 | TMD | S2a |  |  | 0 | 0 |
| S2c |  | 1,700 | TMD-CNST | S2b |  |  | 1 | 0.6×10<sup>6</sup> |
| S2d |  | 10,000 | CG-EQ | S2b | 68,211 | 18×18×22 | 11 | 1.1×10<sup>6</sup> |
| S3a | MuTMC2<br>POPC /<br>POPE /<br>POPS | 100 | EQ | - | 413,104 | 17×17×15 | 0 | 0 |
| S3b |  | 20 | TMD | S3a |  |  | 0 | 0 |
| S3c |  | 1,500 | TMD-CNST | S3b |  |  | 0 | 0 |
| S3d |  | 10,000 | CG-EQ | S3b | 68,391 | 18×18×22 | 6 | 0.6×10<sup>6</sup> |
| S4a | MuTMC2<br>POPC /<br>POPE /<br>POPS /<br>10CHO | 100 | EQ | - | 405,586 | 17×17×15 | 0 | 0 |
| S4b |  | 20 | TMD | S4a |  |  | 0 | 0 |
| S4c |  | 1,700 | TMD-CNST | S4b |  |  | 2 | 1.2×10<sup>6</sup> |
| S4d |  | 10,000 | CG-EQ | S4b | 68,030 | 18×18×22 | 16 | 1.6×10<sup>6</sup> |

